# Synchrotron XRF imaging reveals manganese accumulation in the Golgi and post-synapses of neurons and enhanced uptake in astrocytes

**DOI:** 10.1101/2025.09.05.674493

**Authors:** Ines Kelkoul, Aiyarin Kittilukkana, Luis C.C. Huarte, Hiram Castillo-Michel, Murielle Salome, Stéphane Roudeau, Pauline Belzanne, Matthieu Sainlos, Noémie Pied, Monica Fernandez-Monreal, Daniel Choquet, Richard Ortega, Asuncion Carmona

## Abstract

Manganese is an essential trace metal for humans, but excessive exposure can cause neurotoxicity, including parkinsonian syndromes, cognitive deficits, and has also been implicated in the pathogenesis of neurodegenerative diseases. Globally, tens of millions of people are exposed to elevated manganese levels through drinking water, exceeding the WHO recommended guideline. Despite its public health significance, the cellular and subcellular mechanisms underlying manganese neurotoxicity remain poorly defined, particularly its distribution across brain cell types and its specific intracellular targets. In this study, we investigated manganese accumulation in primary cultures of hippocampal neurons and astrocytes. Using a correlative imaging approach that combined cryo-fluorescence light microscopy with synchrotron X-ray fluorescence imaging, we mapped and quantified manganese at subcellular resolution. Our analysis revealed that manganese preferentially accumulates in the Golgi apparatus of both neurons and astrocytes. In neurons, manganese was also detected in dendrites at the postsynaptic density, suggesting a role in synaptic vulnerability. Quantitative elemental analysis showed that astrocytes accumulated 3 times more manganese than neurons. Furthermore, when neurons were co-cultured with astrocytes, their manganese uptake was significantly reduced, indicating a possible protective or buffering role of astrocytes. These findings identify key cellular and subcellular targets of manganese and highlight the Golgi apparatus as a major regulator of manganese neurotoxicity. This work provides a foundation for understanding cell type–specific responses to manganese exposure and may inform the development of targeted neuroprotection strategies.

**Significance statement:** Manganese neurotoxicity impact millions of people worldwide. Understanding how manganese distributes within brain cells is critical for addressing its neurotoxic effects. This study identifies the Golgi apparatus and postsynaptic density as key sites of manganese accumulation and reveals astrocytes as major regulators of neuronal manganese uptake. These findings provide a cellular framework for developing targeted strategies to mitigate manganese-induced neurotoxicity.

## 1. Introduction

Manganese (Mn) is an essential element required by the body to perform its physiological functions properly. It is involved in the metabolism of many proteins and lipids, as well as the detoxification of reactive species. It also plays a role in energy metabolism and glucose regulation (Li, 2018; Martins, 2025). In the central nervous system, Mn acts as a cofactor for specific enzymes, such as glutamine synthetase. However, Mn is also a toxic element that primarily affects the central nervous system (Ortega, 2025). The balance between essentiality and toxicity is narrow (Vollet, 2016) and overexposure to Mn can lead to various neurological disorders, including motor dysfunction and cognitive impairment (Ortega, 2025). Chronic exposure to high levels of Mn is associated with the development of ‘manganism’, a parkinsonian syndrome whose symptoms resemble those of Parkinson’s disease and include cognitive impairment (Aschner, 2007; Chang, 2010). Mn may also increase the risk of developing idiopathic Parkinson’s disease (Kwakye, 2015). Genetically, mutations in the solute carrier proteins SLC30A10 and SLC39A14, which are involved in Mn transport, have been identified in individuals with familial parkinsonism, leading to severe motor symptoms and mild cognitive impairment (Budinger 2021). Furthermore, various epidemiological studies support the notion that prolonged exposure to Mn can result in multiple cognitive impairments in children (Wasserman, 2006; Bjørklund, 2017; Frisbie, 2019; Kullar, 2019; Mitchell, 2020; Mitchell, 2021). Tens of millions of people worldwide are at risk of being exposed to excessive levels of Mn through their drinking water (Frisbie, 2012). Recently, ambient Mn exposure was associated with cortical brain atrophy in the general adult population (Woo, 2023).

Following overexposure, Mn accumulates in the basal ganglia, particularly in the globus pallidus, but is also observed in other brain regions, such as the choroid plexus, substantia nigra, striatum, frontal cortex, cerebellum, and hippocampus (Ortega, 2025). At the cellular level, studies show that mitochondrial function and microglia-mediated neuroinflammation are affected after Mn exposure (Guo, 2022). Mn also induces the dysregulation of autophagy and exocytosis, as well as causing oxidative stress, endoplasmic reticulum stress and apoptosis, and interfering with neurotransmitter metabolism (Nyarko, 2020; Pajarillo, 2021; Tinkov, 2021). Mn affects mitochondrial morphology and function in both astrocytes (Sarkar, 2018) and neurons (Morcillo, 2021). In neurons, Mn exposure significantly affects dendritic length, spine density, and the number of dendritic endings (Stansfield, 2014; Madison, 2021). Mn targets cell signaling pathways and interferes with neurotransmitter systems, such as the dopaminergic, cholinergic, glutamatergic and GABAergic systems, through multiple mechanisms, including direct interaction with neurotransmitter receptors (Tinkov, 2021; Carmona, 2021). However, the subcellular distribution of Mn in neurons still needs to be identified.

Astrocytes play an active and crucial role in Mn homeostasis in the central nervous system, mediating both protection and neurotoxicity (Sidoryk, 2013; Li, 2024). It is assumed that astrocytes accumulate Mn^2+^ ions through DMT1 (divalent metal transporter 1) and/or by Mn^3+^ binding to the transferrin receptor. In astrocytes, Mn is required for the activity of glutamine synthetase, which catalyzes the conversion of glutamate and ammonium ions into glutamine. Glutamine synthetase contains eight Mn atoms per octamer and accounts for 80% of the total Mn in the brain (Soto, 2021). Astrocytes may accumulate Mn preferentially to neurons, 10 to 50 times more (Sidoryk, 2013; Li, 2024), although quantitative data, such as exact intracellular concentrations in each cell type under identical conditions have not been reported. When present in excess, Mn can disrupt the glutamate/GABA-glutamine shuttle, leading to neuronal damage and contributing to the neurodegenerative process (Li, 2021).

We used synchrotron X-ray fluorescence (SXRF) quantitative imaging to compare the distribution of Mn in single primary rat hippocampal neurons and astrocytes in the same co-culture, which had been exposed to a sub-cytotoxic concentration of Mn. Additionally, using a correlative protocol for imaging metals and proteins (Ortega, 2022; Ortega, 2024), we have elucidated the subcellular distribution of Mn in neurons and astrocytes. This information is key to evaluating the potential protective role of astrocytes, understanding neurotoxicity mechanisms, highlighting subcellular targets, as well as proposing detoxification strategies.

## 2. Materials and Methods

### 2.1 Culture of primary neurons and astrocytes for SXRF imaging

The specificity of our approach lies in the fact that the neurons must be cultured on SXRF-compatible substrates. To this end, we employed silicon nitride membranes (Si_3_N_4_) with a thickness of 500 nm (SiRN-5.0-200-1.5-500, Silson Ltd.). Following the protocol published previously (Perrin, 2015; Domart, 2020; Carmona, 2022; Ortega, 2022) as adapted from Kaech & Banker’s method for culturing hippocampal neurons from embryonic rats (Kaech, 2006). In our protocol, Si_3_N_4_ membranes were sterilized by immersion in absolute ethanol for 2 hours. Any remaining ethanol was removed by washing the membranes twice with sterile, ultra-trace elemental analysis-grade water (Fisher Scientific, France), after which the membranes were dried for 2 hours under a laminar flow hood. To allow cell adhesion, the membranes were coated with 1 mg/mL poly-L-lysine (PLL, P2636, Sigma-Aldrich) in 0.1 M borate buffer (B6768, Sigma-Aldrich) for 2.5 hours at 37 °C. The membranes were then washed twice with sterile ultra-trace elemental analysis grade water and covered with neurobasal culture medium (Gibco, 12349015) to prevent PLL drying and crystal formation.

We recently published a procedure for culturing neurons on Si_3_N_4_ membranes for correlative purposes (Kelkoul, 2025). Here, we describe the specifics of each type of labelling and cell seeding in our study. Primary hippocampal neurons and astrocytes from 18-day-old (E18) Sprague-Dawley rats embryos were seeded and cultured in three different ways according to the experimental objectives, known as the ‘neuron condition’, ‘mixed condition’ and ‘neuron PSD (post-synaptic density) condition’, as explained below. Experimental procedures were in accordance with the European Guide for the Care of Animals used for Experimental and Other Scientific Purposes and the animal care guidelines of the Research Ethics Committee of the University of Bordeaux.

For the culture condition called ‘neuron condition’, dissociated neurons were seeded onto the PLL-coated Si_3_N_4_ membranes at a homogeneous density of 14,000 cells/cm^2^ using equilibrated neurobasal medium supplemented with 0.5 mM GluTmAX (Gibco, 35050038) and B-27 Plus (Gibco, A3582801). This completed culture medium is called NBB27. Two hours after seeding (minimum time to ensure cell adhesion), the membranes were gently transferred to a Petri dish (60 mm diameter) containing an astrocyte feeder layer with 5 mL NBB27 medium and returned to the incubator. At 3 days *in vitro* (DIV3), 2 µM AraC (cytosine α-D-arabinofuranoside hydrochloride, Sigma, C1768) was added to the medium to stop astrocyte cell proliferation in the petri dish. Neurons were cultured until DIV15 and once per week, 1 mL of equilibrated NBB27 was added to compensate for evaporation. In this ‘neuron condition’, only neurons are present on the Si_3_N_4_ membranes.

For the culture condition called ‘mixed condition’, primary rat hippocampal neurons and astrocytes were seeded together on Si_3_N_4_ membranes. For this purpose, the PLL coated Si_3_N_4_ membranes were placed in a coated Petri dish (60 mm diameter) containing 5 mL of NBB27 and 1.5% horse serum. In this case 14,000 cells/cm^2^ were homogeneously seeded onto the Petri dish containing the membranes and immediately returned to the incubator to allow astrocytes and neurons to develop. This ensured both cell types would adhere and grow on the Si_3_N_4_ membranes. At DIV3, the medium was replaced with equilibrated NBB27 without serum. Neurons and astrocytes were cultured until DIV15 and once per week, 1 mL of equilibrated NBB27 was added to compensate for evaporation. In this condition, the Si_3_N_4_ membranes only contain neurons and astrocytes.

In order to image Mn in dendritic spines of the ‘neuron PSD condition’, neurons must be cultured for at least 15 days in close contact with an astrocyte feeder layer. To identify the postsynaptic density, the neurons were transfected with a plasmid coding for the Xph20-eGFP monobody at DIV0 to fluorescently label the postsynaptic protein PSD95 (Rimbault, 2021; Rimbault, 2024). The transfected neurons were seeded at a density of 40,000 cells/cm^2^ onto PLL-coated silicon nitride membranes. Two hours after seeding, the membranes were transferred to a Petri dish containing an astrocyte feeder layer, then returned to the incubator. At DIV3, 2 µM AraC was added to stop astrocyte proliferation. The neurons were cultured until DIV21 to allow maturation and dendritic spine formation, with 1 mL of equilibrated NBB27 being added once per week to compensate for evaporation. In this condition, the Si_3_N_4_ membranes only contain neurons, while the feeder layer of astrocytes is grown on the Petri dish.

### 2.2 Neuron viability assay

Primary rat hippocampal neurons were dissociated and cultured according to the protocol described by Kaech and Banker (Kaech 2006). Viability assays were performed in neurons cultured on coverslips. Dissociated hippocampal neurons were seeded at a density of 200,000 cells in a 60 mm dish containing four PLL-coated 18mm coverslips. Two hours after seeding, the coverslips were transferred to a Petri dish containing a confluent astrocyte feeder layer and 5 mL of NBB27 medium. At DIV3, 2 µM AraC was added to the medium to stop astrocyte proliferation in the Petri dish and neurons were cultured until DIV14. For metal exposure, the coverslips were transferred to a 12-well plates containing 1 mL of culture medium from the dish with the appropriate concentration of Mn. Metal exposure was performed using the medium from the astrocyte feeder layer dish. Neurons were exposed to ultra-pure Mn chloride (Alfa Aesar, 044442.06) for 24h until DIV15, at different concentrations.

Neuronal viability after 24 hours of Mn exposure was determined using the ReadyProbes™ Cell Viability Imaging Kit, Blue/Green (Invitrogen^TM^, R37609). This assay is based on the differential labeling of the nucleus according to the integrity of the plasma membrane. The NucBlue Live reagent (Hoechst 33342) stains the nuclei of all cells (live and dead), while the NucGreen Dead reagent stains only the nuclei of dead cells. The manufacturer’s protocol was adjusted to obtain optimal staining in both channels. Briefly, coverslips were placed in 12-well plates and the medium was replaced with 1 mL of Tyrode’s solution (305 mOsm/L) just before adding 1 drop of NucBlue Live reagent and 2 drops of NucGreen Dead reagent for 45 min incubation at 37°C and 5% CO_2_. This solution was then washed and replaced with 1 mL of Tyrode’s solution to remove fluorescent dyes in the solution prior to microscopy.

Microscopy was performed using a commercial video-microscope, Leica DMI8 inverted (Leica, Germany), with a dry objective 10x (NA 0.32) and equipped with a control in temperature (37°C) and CO_2_. A CoolLED pE-4000 illumination system (CoolLED, US) was used at 405 nm and 490 nm wavelengths and an emission quad band filter DAPI/FITC/TRITC/CY5 was selected to avoid mechanical drift between the imaging of the two dyes. Images were acquired with a Flash 4.0 v2 sCMOS camera (Hamamatsu, Japan) at a 50 ms exposure time for both GFP and DAPI channels. Three replicates of each of the 8 concentrations were imaged and for each coverslip, 7 to 10 different fields of view (1,331×1,331 µm^2^) were acquired throughout the entire coverslip.

### 2.3 Fluorescent labeling and Mn exposure

We only employed fluorescent probes optimized for live-cell microscopy to circumvent the limitations of chemical fixation, which may introduce artefacts that compromise the cellular integrity and is incompatible with elemental analysis (Hansson, 2008; Hackett, 2011; Perrin, 2015; Punshon, 2015; Zohdi, 2015; Jin, 2017; Pushie, 2022). The fluorescence signal remains well preserved after cryo-fixation, thereby enabling efficient localization of the specific subcellular structures of interest by cryo-FM and prior to SXRF imaging (Kelkoul, 2025).

For the ‘neuron condition’, Si_3_N_4_ membranes contained only neurons, and astrocytes were grown in the Petri dish as feeder layer. The neurons were transduced with CellLight™ Golgi-GFP (ThermoFisher Scientific, C10592) according to the manufacturer’s protocol, considering a particle number of 50 per cell. At the time of transduction, the Si_3_N_4_ membranes were transferred to a 12-well dish containing 1 ml of running medium and the required amount of CellLight™ Golgi-GFP. The membranes were then incubated overnight for 16 hours. The following morning, the membranes were transferred to the original Petri dish containing the astrocyte feeder layer. They were then incubated for three days without Golgi staining to increase the transduction rate. After that, a fresh stock solution of Mn at 2 mM was prepared by vortexing for three minutes and sterilized by filtration with 0.22 µm syringe filters. The Si_3_N_4_ membranes were then placed in a new Petri dish containing 2.5 mL of the running medium and 2.5 mL of Mn to reach a final concentration of 250 µM (IC_10_, the 10% viability inhibitory concentration). Thus, in the ‘neuron condition’, the neurons were exposed to Mn in the absence of astrocytes. Following Mn exposure and prior to cryofixation, the membranes were transferred to a 12-well dish and incubated with 5 µg/mL of the nuclear stain Hoechst 33342 (Sigma-Aldrich, B1155) in equilibrated medium at 37 °C for 10 minutes. For the ‘mixed condition’, the transduction and labelling procedure were similar, but in this case Si_3_N_4_ membranes contained both cell types, astrocytes and neurons in co-culture and remained in the astrocyte feeder layer during Mn exposure.

For the ‘neuron PSD condition’, neurons were initially transfected with Xph20-eGFP and cultured until DIV21 to allow for neuronal maturation and the formation of a dense network of dendritic spines. At DIV20, a fresh stock solution of Mn at a concentration of 2 mM was prepared using equilibrated culture medium. This solution was then vortexed for three minutes and sterilized by filtration. The required amount of Mn was then added to the Petri dish to achieve a final concentration of 250 µM. Just before sample preparation by cryofixation at DIV21, the neurons were labelled with SiR-tubulin (silicon rhodamine tubulin, Spirochrome, SC002) to fluorescently label microtubules. To minimize the amount of reagent, the Si_3_N_4_ membranes were transferred into a 12-well dish containing 0.5 ml of equilibrated culture medium and 1 µM of SiR-tubulin. The neurons were then cultured for 1 h at 37 °C (Domart, 2020).

### 2.4 Sample cryo-processing for SXRF analysis

To preserve the cell structure and chemical distribution, the samples were cryopreserved in accordance with the methods previously published (Carmona 2022, Ortega 2024, and Kelkoul 2025). Briefly, after labelling and Mn exposure, the samples were plunged into liquid ethane cooled to -140°C with liquid nitrogen. This process was carried out using an automated vitrification system (Vitrobot Mark IV) from FEI (Thermo Fisher, USA). Immediately prior to vitrification, the membranes were gently and rapidly immersed in a warmed ammonium acetate buffer solution (adjusted to a pH of 7.4 and a concentration of 240 mOsm, the same as the culture medium and prepared using ultra-trace grade water) to remove the extracellular inorganic salts present in the culture medium. Shortly after washing, the samples were placed in the Vitrobot system, where the excess ammonium acetate buffer solution was automatically blotted (blotting parameters: 3 seconds, force 2, 2 blots) using ashless, ultra-absorbent filter paper (Ted Pella, 47000, 595). The membranes were then plunged into chilled liquid ethane to form a thin layer of ice compatible with cryo-FLM (Fluorescence Light Microscopy). After cryofixation, the samples were stored in cryogenic tubes in liquid nitrogen.

After imaging by cryo-FLM (see the next section), the samples were freeze-dried using a Christ Alpha 2-4 LD Plus freeze dryer at -75 °C for 48 hours under primary vacuum (2·10^−3^ mbar). The freeze-dried samples were then allowed to gradually reach room temperature under vacuum conditions for 5 hours. Once room temperature was reached, the vacuum was released and dry air from desiccant silica beads was allowed to enter. Finally, the freeze-dried samples were stored in a desiccant cabinet at room temperature until SXRF analysis.

### 2.5 Cryo-fluorescence light microscopy (cryo-FLM)

The cryo-fixed membranes were transferred to a Leica (Germany) commercial wide field EM-Cryo-CLEM microscope at the Bordeaux Imaging Centre (BIC). This system keeps the samples in liquid nitrogen vapor throughout the entire process, including transfer, storage and observation. Cryo-FLM images were acquired using Metamorph software (Molecular Devices). The physical length of the images was set to 2,304 pixels at 104 nm/pixel using an HC PL APO 50x/0.90 NA cryo objective and an Orca-Fusion BT camera (Hamamatsu Photonics France SARL).

Golgi and Hoechst 33342 imaging was performed using a GFP filter (470/40 nm excitation, BP 525/550) and a DAPI filter (360/40 nm excitation, BP 470/40 nm), respectively. The same GFP filter and a Y5 filter (620/60 nm excitation, BP 700/75 nm) were used for GFP-xph20 and SiR-tubulin imaging. For each wavelength, a z-stack with a thickness of 3–4 µm was acquired in order to image the full cell volume. Bright-field images were also acquired for the same regions to improve final correlation.

After freeze-drying the samples, the same neurons were imaged again at room temperature using the DAPI filter and brightfield illumination to map the entire membrane and facilitate identification of the regions of interest during SXRF imaging. See Supplementary Material 1 for details of the correlative protocol.

### 2.6 Synchrotron X-ray fluorescence imaging

SXRF imaging experiments were performed at the ID21 beamline of the ESRF (European Synchrotron Radiation Facility) using two experimental setups. In the first setup, a monochromatic beam of 7.4 keV was focused to 890 nm x 230 nm, providing a flux of 1.1 × 10^11^ photons/s. Samples were raster scanned in continuous mode with an integration time of 100 ms/pixel and a step size of 500 nm/pixel. The larger spatial resolution enables a significant number of whole cells to be analyzed for each condition and cell type, providing statistically significant information (Salome, 2013). In the second setup, the recently commissioned nanoscope at ID21 was used for imaging dendritic spines at room temperature and under vacuum conditions. This new setup enables a 10.2 keV monochromatic beam to be focused down to a size of 167 nm x 151 nm, providing a flux of 9.9 x 10^10^ photons/s. Samples were raster scanned continuously with an integration time of 100 ms/pixel and a step size of 200 nm/pixel.

The XRF signal was recorded using X-ray detectors based on multi-element silicon drift diodes, which were positioned at 90° to the incident beam. The summed X-ray fluorescence spectra from these detectors were fitted using Python scripts and the PyMCA library (Solé, 2007), and were calibrated against the signal derived from a thin-film X-ray fluorescence standard (AXO Dresden GmbH, Germany). The resulting areal mass density distributions of the detected chemical elements were saved individually as 32-bit TIFF images.

For element quantification, two regions of interest (ROIs) were defined in the quantitative images using FIJI software (Schindelin et al., 2012): one ROI from the neuron and one for the background. The intracellular element content, expressed in ng/mm^2^, was calculated as the difference between the mean intensity of the two ROIs (see Supplementary Material 2). These values were then converted to micromolar units (see Supplementary Material 3) by measuring the thickness of living neurons and astrocytes using a calibrated Z microscope (see Supplementary Material 4). The elemental content was then compared across two conditions: neurons exposed to Mn in the absence of astrocytes (the ‘neuron’ condition); neurons and astrocytes exposed to Mn in co-culture (the ‘mixed’ condition).

### 2.7. Correlative imaging

For correlative imaging purposes, the Icy software (de Chaumont, 2012) version 2.4.3 (http://icy.bioimageanalysis.org/) and the eC-CLEM plugin (Paul-Gilloteaux, 2017) were used. Cryo-FLM images were presented as the maximum projection of the acquired z-stack to enable superposition with XRF maps. The procedure for correlating cryo-FLM and SXRF images is outlined in Supplementary material 1. Three groups of images were superimposed. The first group comprised the cryo-fluorescence images (Golgi, nucleus and bright field); the second comprised the images of freeze-dried samples (bright field, nucleus and Golgi); and the third comprised the SXRF images of the chemical elements.

### 2.8 Statistics

The statistical analysis of the cell viability assays was performed in two steps. First, the microscopy images were analyzed using an ImageJ macro to segment the cells in the two measurement channels: total cells and dead cells. These results were then processed using Rstudio Notebooks to perform statistical analysis, plot dose-response curves and calculate IC_10_ and IC_50_ values (viability inhibitory concentration at 10% and 50%). Three dose-response models were empolyed, and the weighted mean was calculated to plot the final dose-response curve and determine de IC values. R software (version 4.3.0) and Rsudio (version 2023.12.0), Tidyverse (2.0.0) (Wickham, 2019) and drc (3.0-1) (Ritz, 2015) R packages were used.

GraphPad Prism 8.4.3 for Windows (Boston, Massachusetts, USA; www.graphpad.com) was used for the statistical analysis of SXRF data to produce box plots and the associated statistics (see Supplementary materials 5 for details). The number of analyses in each group was n = 23, 15 and 19 for ‘neuron condition’, neurons ‘mixed condition’ and astrocytes ‘mixed condition’, respectively. The Shapiro-Wilk test showed that the data were not normally distributed, so Mann-Whitney non-parametric tests were performed to compare the groups two by two.

## 3. Results

### 3.1 Manganese subcellular imaging in neurons and astrocytes

As described in Section 2.3, we employ fluorescent probes optimized for live-cell labeling to circumvent the limitations of chemical fixation. The location of the fluorescent markers of interest was imaged by cryo-FLM and their positions recorded. Subsequently, the same cells were analyzed using SXRF to visualize the distribution of chemical elements (Fig. 1).

**Fig. 1.**
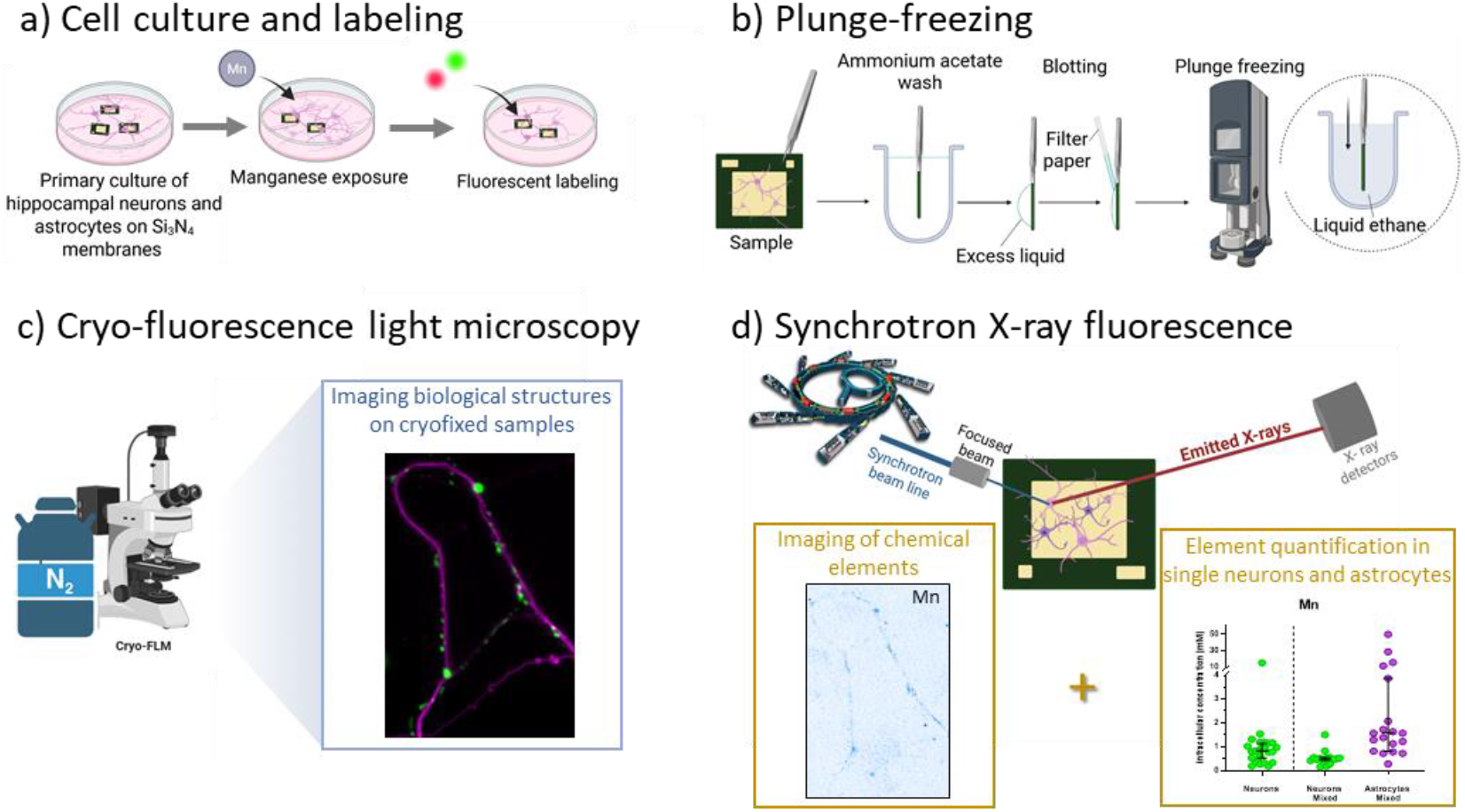
Schematic overview of the sample preparation and correlative imaging workflow combining cryo-FLM and synchrotron XRF imaging. a) Primary cultures of hippocampal neurons and astrocytes onto silicon nitride membranes, exposed to Mn and fluorescent labelled. b) After labeling, samples were rapidly plunge-frozen in liquid ethane using an automated device. c) Cryo-FLM was performed to visualize the distribution of fluorescently labeled cellular structures on cryofixed samples. d) The same regions of interest were analyzed by SXRF for both correlative imaging purposes and element quantification.

In control cells, not exposed to Mn, the quantification of this element was below the detection limit and could not be imaged in astrocytes or neurons (see Supplementary material 6). An example of the elemental distribution in a cultured astrocyte from the ‘mixed condition’ exposed to 250 µM of Mn for 24h (corresponding to IC_10_, see supplementary material 7) is shown in Fig. 2; other examples can be found in Supplementary material 8. The cryo-FLM image (Fig. 2a) shows the Golgi apparatus located in a compact cytoplasmic area close to the nucleus. The elemental distribution (Fig. 2b) obtained by SXRF imaging shows that P and K are distributed throughout the cell, whereas Ca and Mn are located in a compact area of the cytoplasm. Merging the elemental distributions (Fig. 2c) reveals that Mn and Ca co-localize perfectly and that Mn accumulation occurs adjacent to the phosphorous-rich region of the nucleus. Merging cryo-FLM and SXRF images (see section 2.7 and Supplementary material 1), allows obtaining the distribution of the chemical elements and organelles to be visualized in the same cell. The Mn distribution (red) partially co-localizes with the green fluorescence of the Golgi apparatus (Fig. 2d). This partial co-localization could be explained by differences in protein localization at the sub-compartment level of the Golgi apparatus (see Discussion and Conclusion).

**Fig. 2.**
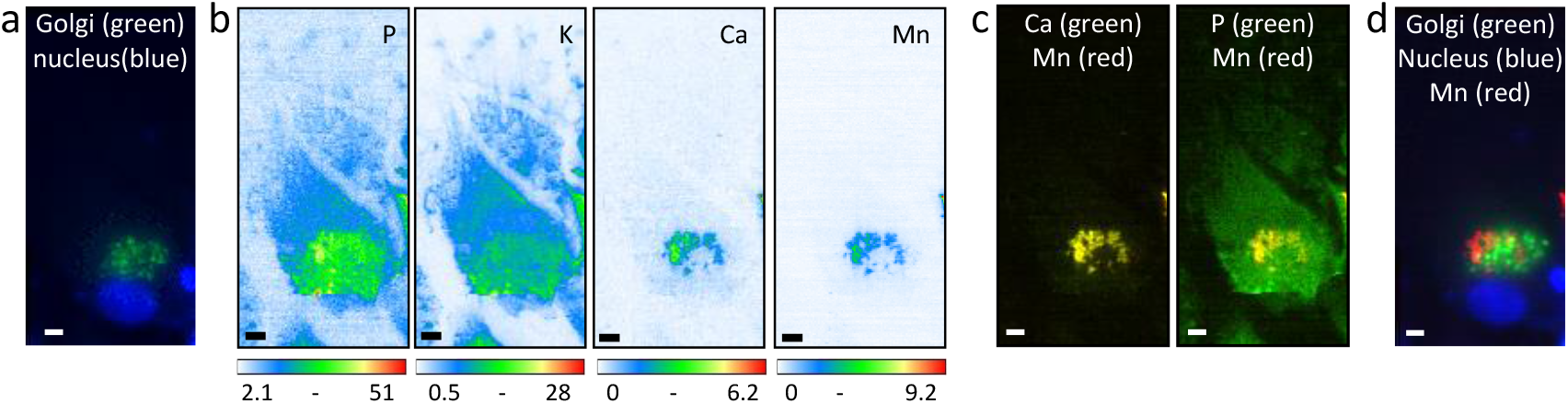
Organelle and elemental imaging in astrocytes after Mn exposure. a) Merged images of the nucleus (blue) and the Golgi apparatus (green) obtained by cryo-FLM. b) Elemental distribution of P, K, Ca and Mn obtained by SXRF, scan size 48 µm x 95 µm, step 0.5 µm/pixel, scan time 100 ms/pixel. The color scales represent the quantitative values per pixel, expressed in ng/mm^2^. c) Merged images of Ca (green) and Mn (red), showing a perfect colocalization of both elements (yellow) and of P and Mn showing Mn is restricted adjacent to the P-rich nucleus. d) Merged images of Mn (red), Golgi apparatus (green) and nucleus (blue). Scale bars: 5 µm.

Figure 3 shows an example of the elemental distribution in a primary hippocampal neuron exposed to Mn in the presence of astrocytes (‘mixed condition’). Other examples can be found in Supplementary Material 9. The cryo-FLM images (Fig. 3b) reveal a dotted pattern of the Golgi apparatus on the right side of the nucleus. The elemental distribution (Fig. 3a) shows that phosphorus (P) and potassium (K) are distributed throughout the cell, while calcium (Ca) and Mn follow a dotted pattern and are principally located on the right side of the cell. Superimposing the cryo-FLM and SXRF images (Fig. 3c) reveals that Mn is located in the Golgi apparatus region, although the individual vesicles do not show strict colocalization. Superposition of Mn and P shows that Mn is restricted to a cytoplasmic region, while superposition of Ca and Mn shows that both elements colocalize in the Golgi area (Fig. 3d, white arrow). Interestingly, outside the cytoplasmic region, some Mn vesicles are found along the neuronal ramifications (magenta arrows in Figs 3a and 3d). Similarly to the results found for astrocytes, Mn partially colocalizes with the green fluorescence of the Golgi region, and perfect colocalization occurs between Ca and Mn.

**Fig. 3.**
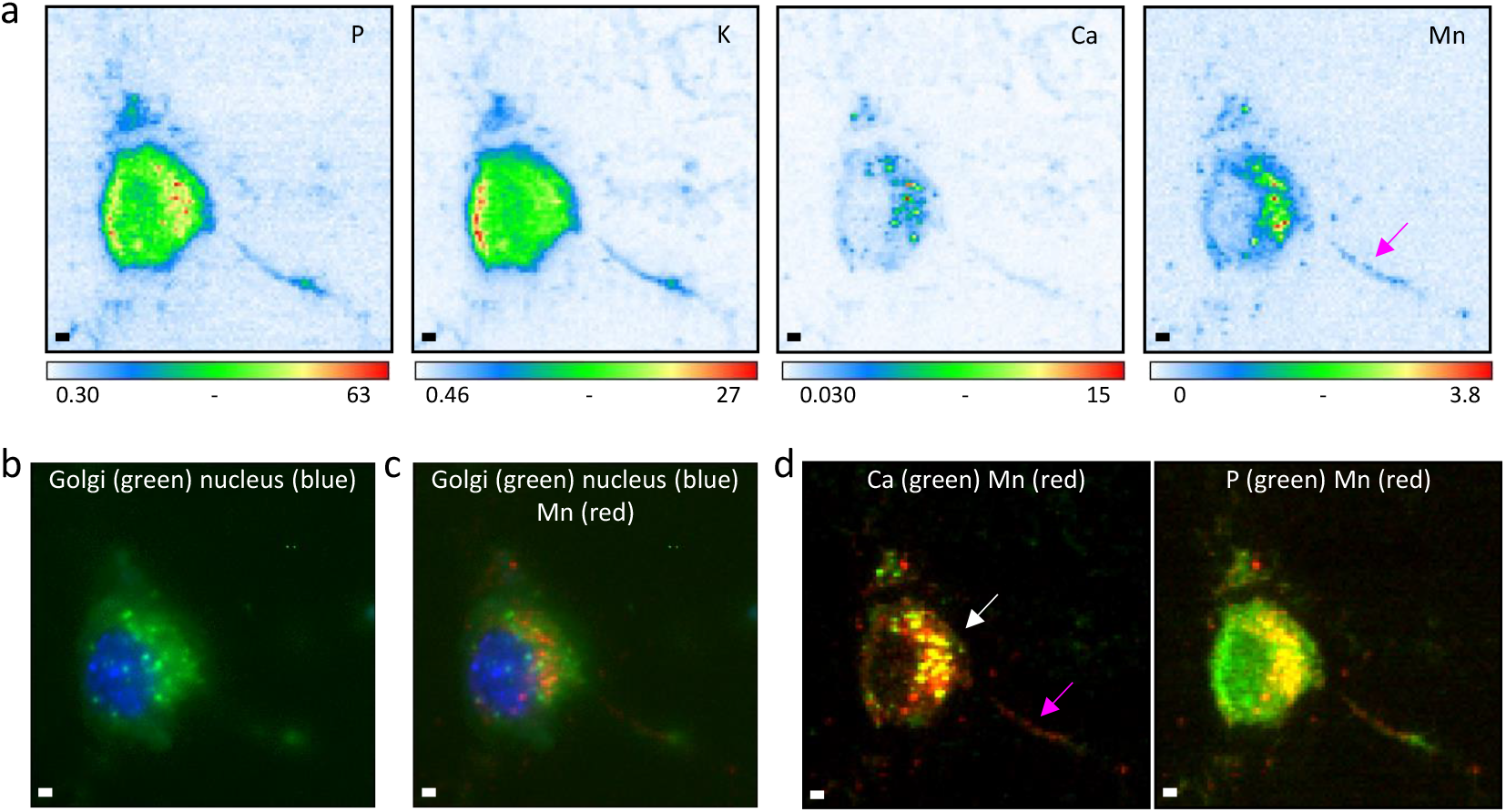
Organelle and elemental imaging in primary neurons after Mn exposure in the presence of astrocytes. a) Elemental distribution of P, K, Ca and Mn obtained by SXRF, scan size 50 µm x 50 µm, step 0.5 µm/pixel, scan time 100 ms/pixel. The color scales represent the quantitative values per pixel, expressed in ng/mm^2^. b) Merged images of the nucleus fluorescence (blue) and the Golgi apparatus (green) obtained by cryo-FLM. c) Superimposed images of Mn (red), Golgi (green) and nucleus (blue). d) Merged images of Ca and Mn, showing colocalization of both elements in the Golgi region (yellow, white arrow) and Mn reaching the dendritic ramification (magenta arrow), merged images of P and Mn, showing most of the Mn in a restricted area of the cytoplasm. Scale bars: 5 µm.

Figure 4 shows an example of the elemental distribution in a primary hippocampal neuron exposed to Mn in the absence of astrocytes (‘neuron condition’). Other examples can be found in Supplementary Material 10. The cryo-FLM images (Fig. 4b) show the Golgi apparatus displaying a punctate pattern beneath the nucleus. Fig. 4a shows the elemental distributions, with P and K distributed throughout the cell and Ca and Mn concentrated in areas of the cytoplasm. Superimposing the cryo-FLM and SXRF images (Fig. 4c) reveals that, while Mn is still located in the Golgi region, it does not follow a dotted pattern, as opposed to the images of the ‘mixed condition’, instead exhibiting a more homogeneous distribution (Figures 3a and 4a; Supplementary materials 9 and 10). Superposition of Ca and Mn (Fig. 4d) shows that both elements colocalize in the Golgi region. Superposition of P and Mn confirms that Mn is restricted to a cytoplasmic region (Fig. 4d). Interestingly, outside the cytoplasmic region, some Mn is found along the dendrites, as observed in neurons from both the ‘mixed’ and ‘neuron’ conditions (Figures 3a and 4a, magenta arrows).

**Fig. 4.**
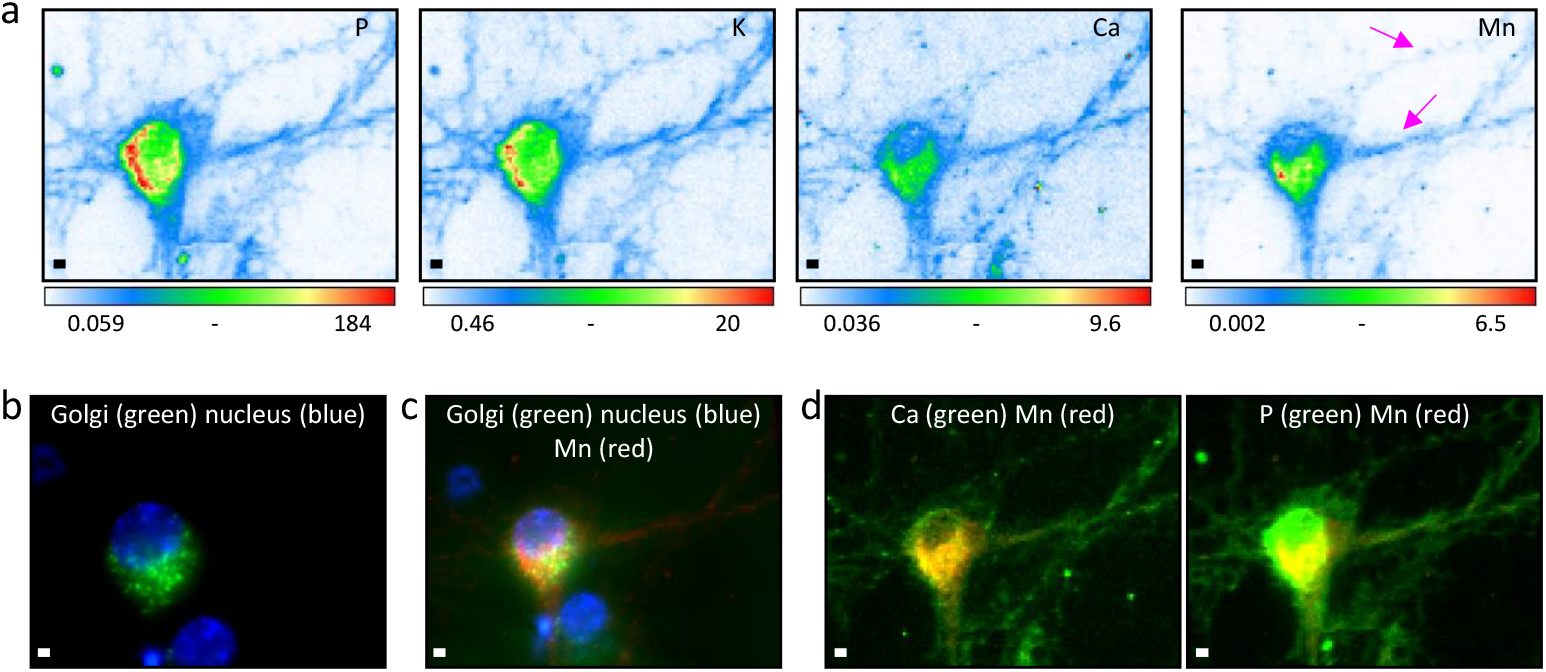
Organelle and elemental imaging in primary neurons after Mn exposure in the absence of astrocytes. a) Elemental distribution of P, K, Ca and Mn obtained by SXRF, scan size 60 µm x 46 µm, step 0.5 µm/pixel, scan time 100 ms/pixel. The color scales represent the quantitative values per pixel, expressed in ng/mm^2^. b) Merged images of the nucleus (blue) and the Golgi apparatus (green) obtained by cryo-fluorescence light microscopy. c) Superimposed images of Mn (red) and Golgi (green) and nucleus (blue). d) merged images of Ca and Mn, showing colocalization of both elements (yellow) and merged images of P and Mn, showing Mn in a restricted area of the neuron. Scale bars: 2 µm.

To investigate whether Mn reaches regions other than the Golgi apparatus in neurons, we imaged dendrites far from the soma (Fig. 5). The P and K distributions (Fig. 5a) reveal the morphology of neuronal branching, with Ca and Mn distributed as puncta (see also Supplementary material 11). Merged images of P and Mn (Fig. 5b and 5c) reveal Mn dots (red) scattered across neuronal branches. In contrast, the merged images of Ca and Mn (Fig. 5b and 5c) reveal two types of Mn occurrence. Some large Mn puncta colocalize with calcium (white arrows), while smaller ones do not (magenta arrows). Whole neuron imaging revealed that Mn localization in neurons is not restricted to the Golgi region, and that Mn-containing vesicles colocalize with calcium vesicles.

**Fig. 5.**
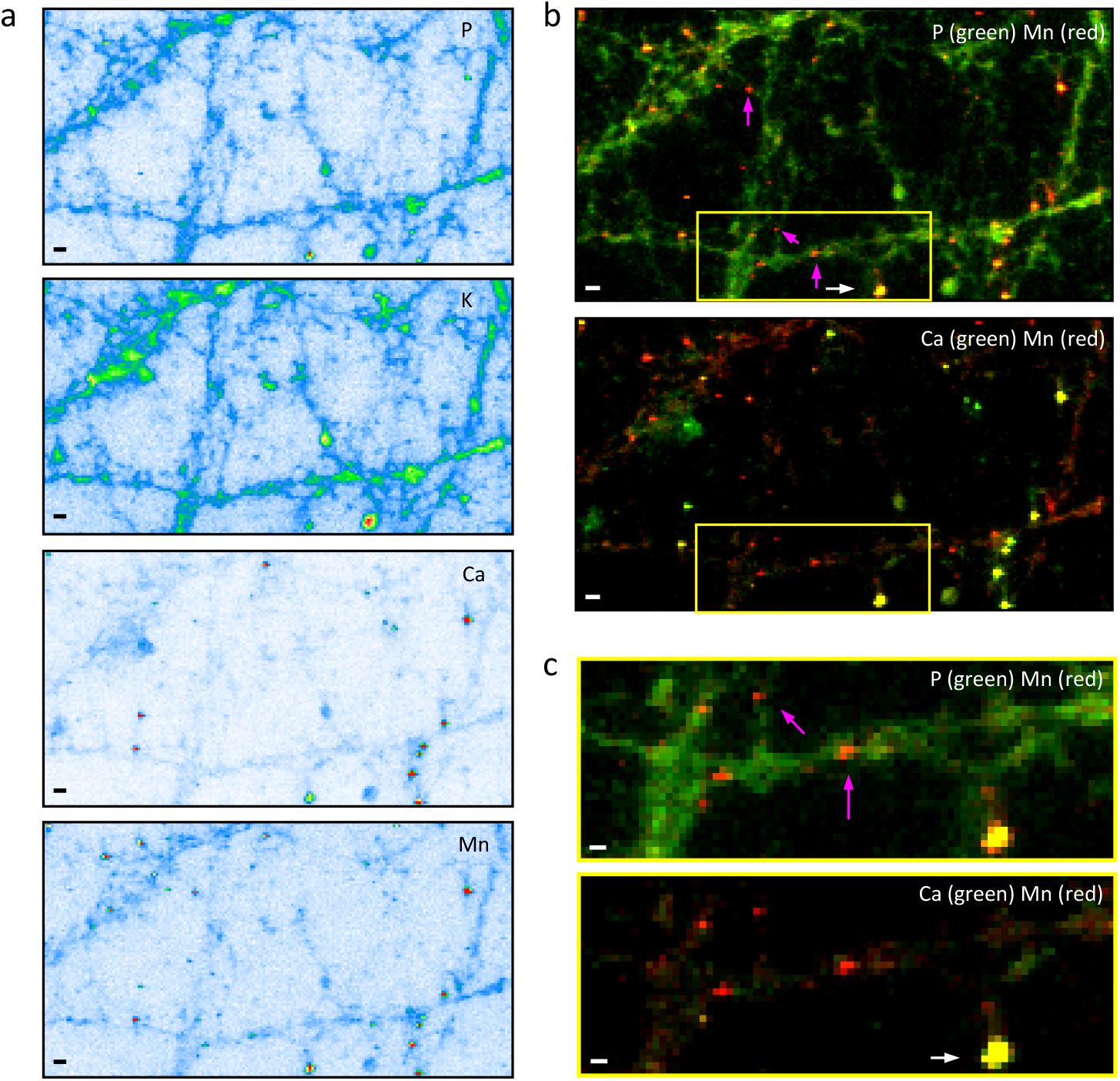
Elemental imaging in the dendritic network of hippocampal neurons. a) Elemental distribution of P, K, Ca and Mn obtained by SXRF, scan size 74 µm x 40 µm, step 0.4 µm/pixel, scan time 100 ms/pixel. b) Merged images of Mn and P, and merged images of Mn and Ca, highlighting the presence of Mn alone in small dots in the neuronal branching (magenta arrows) and colocalizing with Ca in large puncta (white arrows). c) Zoomed region from the white square in b) showing Mn along dendrites and Mn and Ca colocalization. Scale bar 2 µm (a, b); 1µm (c).

### 3.2 High resolution manganese imaging in dendritic spines

To investigate whether Mn reaches dendritic spines, we fluorescently labelled the PSD-95 protein to localize post-synaptic compartments and the tubulin protein to image microtubules (Fig. 6 and Supplementary material 12). Fig. 6a shows an overview of the neuronal network. PSD-95 fluorescence was observed along the microtubules in the selected region of interest. These areas were imaged using cryo-FLM to discern the distribution of tubulin (magenta) and PSD-95 (green) (see Fig. 6b). The same region was analyzed using SXRF imaging to determine the distribution of elements such as Mn and Zn (Fig. 6c). Using ICY and eC-CLEM (Section 2.7), the cryo-FLM and SXRF images were superimposed to reveal the overlap between the distribution of PSD-95 and the localization of Mn and Zn (Figure 6d).

**Fig. 6.**
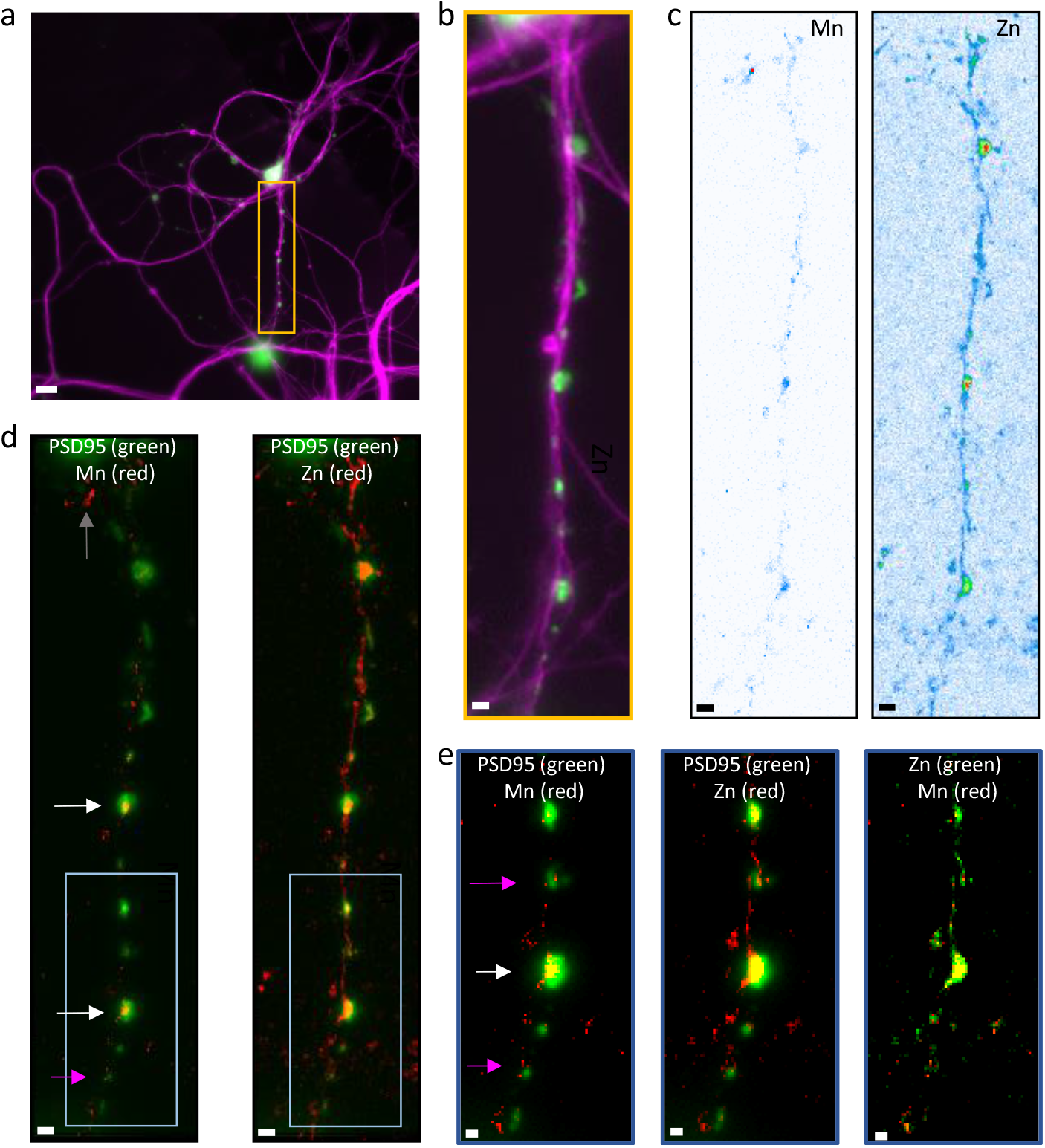
Elemental imaging in dendritic spines of primary neurons exposed to Mn. a) Confocal fluorescence images of tubulin (magenta) and PSD-95 (green) in living primary hippocampal neurons at DIV21. b) Cryo-FLM images of the framed region in a, showing fluorescence of tubulin (magenta) and PSD-95 (green) in immobilized cryogenic conditions. c) Elemental distribution of Zn and Mn obtained by SXRF, scan size 85 µm x 20 µm, step 0.2 µm/pixel, scanning time 100 ms/pixel. d) Superimposed images of Mn (red) and PSD-95 (green) distribution and of Zn (red) and PSD-95 (green) showing Mn reaching the postsynaptic density (white arrows), in the close vicinity of the postsynaptic density (magenta arrows) and along the microtubules (gray arrow). e) zoom in the region framed in d to show Mn and Zn location respectively to PSD-95, and Mn and Zn superposition. Scale bars: 10 µm (a); 2 µm (b, c, d); 1 µm (e).

Three types of Mn distribution can be distinguished from the superimposed images (Fig. 6d and Supplementary material 12d). Firstly, Mn is present in the highly fluorescent PSD-95 spots (Fig. 6d, white arrow, and Supplementary material 12e). Secondly, Mn is found in close proximity to small PSD-95 spots (Fig. 6d, magenta arrows, and Supplementary material 12f). Thirdly, some larger Mn spots are present along microtubules (Fig 6d, grey arrow, and Supplementary material 12g). This is clearly visible in the zoomed regions (Fig. 6e and Supplementary material 12e-g): Mn reaches large postsynaptic compartments (Fig. 6e white arrow), or is located around smaller postsynaptic compartments (Fig. 6e magenta arrows). Zn is also present in the postsynaptic compartments (Fig. 7e and Supplementary material 12e-g), as expected from our previous findings regarding Zn distribution in neurons (Ortega, 2022). Finally, the superposition of Zn and Mn indicates that the two elements are not strictly colocalized, although both elements are located within the larger PSD-95 fluorescent structures, or around the smaller ones (Fig. 7e and Supplementary material 12e-g).

**Fig. 7.**
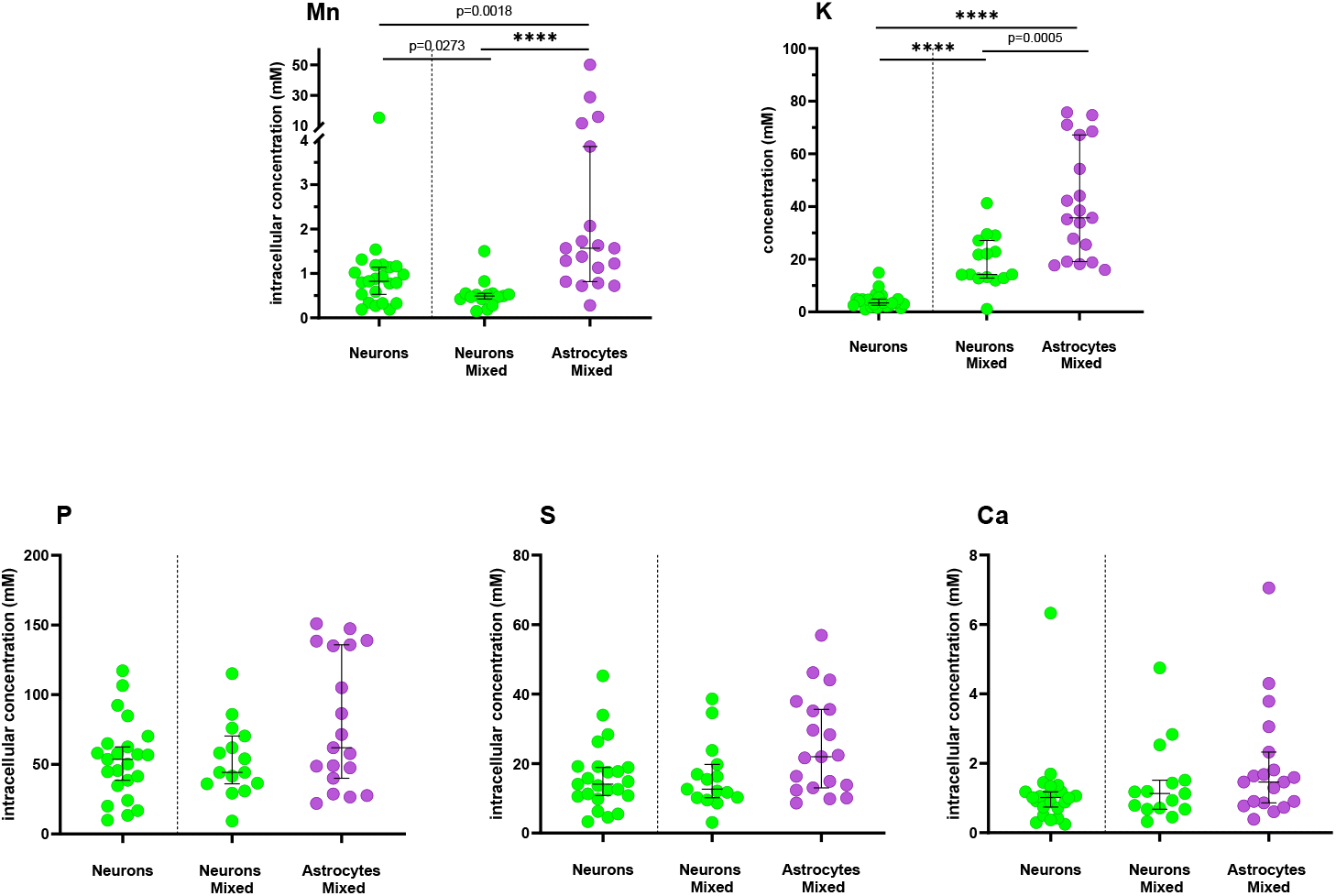
Elemental content in neurons and astrocytes after Mn exposure. Each dot in the scatter-plot represents the quantitative values obtained for one neuron (green dots) or astrocyte (magenta dots). For each condition, the black lines represent the median and the confidence interval at 95%. Data did not fulfil the Shapiro Wilk test and groups were compared using Mann-Whitney test. The number of cells measured for each group was 23, 15 and 19 for neurons, neurons in mixed conditions and astrocytes respectively. p < 0.0001 = ****. For details on statistical data analysis see section 2.8 and Supplementary material 5.

### 3.3 Multi-element quantification in neurons and astrocytes

Two types of cell exposure to Mn were compared. In the ‘neuron condition’, the neurons were cultured alone on the Si_3_N_4_ membranes and in close contact with a feeder layer of astrocytes grown on the Petri dish, except during Mn exposure, which was conducted without the astrocyte layer (see section 2.1). In the ‘mixed condition’, neurons and astrocytes were co-cultured on the Si_3_N_4_ membranes during the entire process, including Mn exposure. Mn exposure was set to 250 µM Mn for 24h, corresponding to the IC_10_ viability assay. Element quantification was compared on three groups: 1) neurons cultured alone from ‘neuron condition’, 2) neurons cultured with astrocytes from ‘mixed condition’ and 3) astrocytes from ‘mixed condition’.

The intracellular content, as calculated by SXRF data analysis is initially expressed in ng/mm^2^. As explained in section 2.6 and Supplementary material 3, these values were transformed into mM units. To do this, the thickness of each cell type, 15.5 ± 2.3 µm for neurons and 5.8 ± 0.8 µm for astrocytes was measured in living cells (Supplementary material 4). Therefore, the expressed concentrations in mM, are volumetric and take into account the volume under hydrated conditions.

With regard to Mn quantification, a significant difference was observed between neurons exposed to Mn in the presence or absence of astrocytes (p = 0.0273). Mn accumulation is almost halved in neurons from the ‘mixed condition’ compared to neurons from the ‘neuron condition’ (see Fig. 7). The median and median absolute deviation for neurons were 490 ± 60 µM and 820 ± 340 µM, respectively (see Supplementary Material 3). Astrocytes have a higher capacity to accumulate Mn than neurons, with a median and median absolute deviation of 1,600 ± 800 µM, as well as high intracellular variability. Mn accumulation can vary considerably among astrocytes, with two potential populations of high and low Mn content.

Element quantification also reveals statistically significant changes in potassium content. There is a lower potassium in both neuronal populations compared to astrocytes (Fig. 7). There is a greater decrease in intra-neuronal potassium in neurons that have not been in contact with astrocytes during exposure. These results confirm that neuronal integrity is compromised when Mn accumulates and that astrocytes protect against toxicity.

Our results also show statistically significant changes in potassium content. A decrease of potassium occurs in both neuronal populations with respect to astrocytes (Fig. 7). Moreover, in the ‘neuron condition’, potassium content in neurons is statistically significantly lower than in neurons from the ‘mixed condition’. This result may indicate that the integrity of neurons is compromised resulting in potassium loss when they accumulate more Mn. The intracellular accumulation of other elements such as P, S and Ca is not statistically different between neurons and astrocytes and is neither modified by the presence or absence of astrocytes layer (Fig. 7).

## Discussion and Conclusion

Mn is an essential element found at trace levels in human tissues, with an estimated total amount of 390 µg in the adult human brain (ICRP23, 1975). This typically corresponds to 0.3 µg/g of wet weight (Williams, 2012) or 1.3 µg/g of dry weight (Ramos, 2014). Due to this low concentration, detecting and imaging Mn in single cells is challenging and requires highly sensitive spatially resolved analytical techniques. Due to such analytical challenges, imaging endogenous levels of Mn (<1 µg/g) in subcellular compartments is below the detection limit of current micro-analytical techniques, as was found in this study when using SXRF (Supplementary material 6). However, Mn subcellular imaging can be achieved on cells supplemented with Mn. Probably due to its essential functions, Mn is toxic only at relatively high extracellular concentrations. In this study, we determined that the IC_10_ for primary rat hippocampal neurons was 250 µM (Supplementary material 7). This value falls within the same concentration range as that obtained for cell lines in previous studies (Cai, 2010; Carmona, 2010; Gandhi, 2018).

After Mn exposure, only a few studies have addressed the subcellular distribution of Mn in brain cells or, more generally, in mammalian cells to date. This has been achieved by experimental approaches with high spatial resolution and high analytical sensitivity, such as synchrotron XRF (Carmona, 2010; Leoni, 2011); or with the recent development of chemical biology tools to image Mn^2+^ ions in living cells (Park, 2022; Kahali, 2024). These fluorescent biomolecular sensors are of great promise for understanding dynamic processes involving Mn^2+^ ions in cells, but they are not yet commercially available and have not yet been used in brain cells. Moreover, they are restricted to imaging labile Mn^2+^ ions and cannot image the total cellular pool of Mn, which includes Mn bound to biomolecules.

Using correlative synchrotron XRF imaging and fluorescence microscopy, we previously demonstrated that the main subcellular site of Mn accumulation in neuron-like PC12 rat cells exposed to Mn was the Golgi apparatus (Carmona 2010, Carmona 2014). Synchrotron XRF imaging of primary mouse hippocampal neurons and of neuron-like N2a mouse cells exposed to Mn showed that Mn was mainly located in perinuclear areas resembling the Golgi apparatus (Daoust 2014), in terms of size, shape and subcellular localization. However, strict identification of the organelles was not performed. A similar result was obtained using synchrotron XRF imaging on primary mouse midbrain neurons exposed to Mn, which showed perinuclear Mn accumulation (Dučić 2013; Dučić 2015). In our study, we imaged the distribution of Mn and the localization of the Golgi apparatus. We report their co-localization in primary rat hippocampal neurons and astrocytes exposed to Mn and demonstrate that the Golgi apparatus is the main organelle of Mn accumulation in these brain cells. This finding is consistent with studies of other cell types. Mn^2+^ ions and total Mn were located within the Golgi apparatus in HEK293T primary human embryonic kidney cells (Das 2019). We have also observed Mn accumulation in the Golgi apparatus of HeLa human cervical carcinoma cells overexpressing a mutant SLC30A10 protein (Carmona 2019). Our results provide new data showing that the Golgi apparatus is the main storage organelle for Mn in neurons and astrocytes (Figs. 2-4 and Supplementary material 8-10), confirming a mechanism that has been observed for other cell types (Carmona 2010, Carmona 2014, Das 2019, Carmona 2019).

The accumulation of Mn in the Golgi apparatus coincides with the localization of the secretory-pathway Ca^2+^-ATPase SCPA1, which is known to transport Ca and Mn ions from the cytosol to the Golgi apparatus (Missiaen 2004). SCPA1 can transport both Ca^2+^ and Mn^2+^ ions and is expressed mainly in the trans-Golgi network (Pizzo 2010). Our results demonstrate that Mn co-localizes with Ca in the Golgi apparatus region (Fig. 2-4 and Supplementary material 8-10). Furthermore, we found that the distribution of Mn/Ca does not correlate strictly with the localization of the CellLight^TM^ Golgi-GFP marker (Fig. 2-4 and Supplementary material 8-10). This can be explained by the cis-Golgi localization of the N-acetylglucosaminyltransferase used in CellLight^TM^ Golgi-GFP construct (Roth 1994), which differs from the trans-Golgi localization of SCPA1. Our results strongly suggest that SPCA1 and the Golgi apparatus play an important role in Mn storage and neuroprotection. This is consistent with the observation that HeLa cell viability increases when SPCA1 activity is increased (Mukhopadhyay 2011). In glial cells, SPCA1 protects against Mn toxicity, increasing cell viability in overexpressing cells and decreasing it in silenced cells (Bhojwani-Cabrera, 2024). At the organ level, magnetic resonance imaging of animals exposed to Mn has shown Mn accumulation in brain regions with high SPCA1 expression, such as the CA3 region of the hippocampus (Sepúlveda, 2012). These results suggest that the Golgi apparatus and SPCA1 play a key role in Mn storage and neuroprotection. However, following exposure to high levels of Mn, this protective mechanism may become overwhelmed, resulting in Mn toxicity and impairment of Golgi apparatus functions (Carmona, 2010; Bhojwani-Cabrera, 2024). For example, high levels of exposure to Mn lead to fragmentation of the Golgi apparatus and loss of the Golgi ribbon structure in neurons and glial cells (Sepúlveda, 2012; Bhojwani-Cabrera, 2024). Golgi fragmentation is a common feature of many neurodegenerative diseases, including Alzheimer’s disease, Huntington’s disease, amyotrophic lateral sclerosis, and Parkinson’s disease (Makhoul, 2019). Therefore, the Golgi apparatus is involved in both neuroprotection, through Mn storage, and neurotoxicity, when its structure begins to deteriorate, resulting in altered Golgi functions and neurodegeneration.

Although most of the Mn reaches the Golgi, other organelles and subcellular regions may also accumulate Mn in smaller amounts, which may help explain the pleiotropic neurotoxic effects of Mn. There are some indications in the literature that Mn can accumulate in neurites. Synchrotron XRF imaging has observed discrete Mn spots along neurites in primary mouse hippocampal neurons (Daoust, 2014) and primary mouse midbrain neurons (Dučić, 2015), although the exact localization of this Mn was not characterized further. Using a correlative metal and protein co-localization approach, our results show that Mn co-localizes with the postsynaptic density in dendrites of primary rat hippocampal neurons (see Figs. 6 and Supplementary material 12). Compared to zinc, which is physiologically enriched in the PSD (Ortega, 2022), Mn displays a different distribution within the PSD (see Figures 6 and Supplementary material 12). While zinc fills the fluorescently labelled PSD region, Mn accumulates in smaller subregions. This indirectly suggests that Mn may target sites other than zinc-binding molecules, such as the Shank scaffolding proteins known to bind zinc in the PSD (Baron, 2006). While the exact mechanism of Mn accumulation within the PSD remains unclear, our results demonstrate that Mn can reach the postsynaptic compartment, potentially triggering toxic effects. This finding is consistent with a recent study demonstrating that exposure to Mn in mice leads to significant Mn accumulation in the hippocampus, resulting in neuronal damage and reduced dendritic spine density (Bi 2024). This is also consistent with electron microscopy observations in mice treated with Mn, which showed decreased electron density in postsynaptic areas and decreased, dispersed synaptic vesicles in presynaptic areas (Villalobos, 2015). In our study, the lack of fluorescent markers for presynaptic compartments that are compatible with live cell imaging meant that Mn accumulation at the presynaptic level could not be studied. However, this accumulation is indirectly suggested by the detrimental effects of Mn on presynaptic mechanisms (Guilarte, 2008).

Our results demonstrate the significant potential of combining SXRF element imaging and protein fluorescence microscopy to identify sites of subcellular metal distribution, which is essential for understanding the mechanisms of metal neurotoxicity (Kelkoul, 2025). Another significant advantage of single-cell SXRF spectro-imaging is the ability to describe and quantify metal distribution in various cell types in co-culture. Using this capability, we quantified the element content in neurons cultured alone or in co-culture with astrocytes, showing that the latter contained approximately 40% less Mn (Fig. 7). This indicates that astrocyte presence reduces neuronal Mn uptake, contributing to neuroprotection. In the absence of astrocytes, we also observed that Mn accumulates excessively in the Golgi region, resulting in a broader and higher local distribution (Figures 3a and 4a; Supplementary materials 9 and 10). This may indicate dysfunction in the storage capacity of the Golgi apparatus, as has been reported in PC12 cells exposed to excess Mn (Carmona, 2010).

The neuroprotective role of astrocytes is also supported by higher potassium levels in neurons co-cultured with astrocytes compared to neurons cultured alone (Fig. 7). Potassium loss is a marker of cytotoxicity that can be mediated by over-activated potassium and/or ionotropic glutamate receptor channels (Yu, 2003). Furthermore, Mn quantification in the ‘mixed condition’ suggests that astrocytes absorb three times more Mn than neurons. The sequestration of Mn by astrocytes may indirectly limit its availability to neurons. Astrocytes account for approximately 80% of total Mn content in the brain under physiological conditions (Soto, 2021), making them the primary pool of Mn. It is often assumed that astrocytes can accumulate 10-to 50-fold more Mn than neurons (Sidoryk, 2013; Li, 2024), although exact intracellular concentrations have not yet been measured in either cell type under identical conditions. Our results indicate that, under identical Mn exposure conditions, astrocytes accumulate approximately three times more Mn than neurons (Fig. 7). However, we observed large interindividual variability in astrocyte Mn content, possibly indicating two types of astrocytes characterized by low and high Mn levels. In astrocytes with the highest Mn levels, the Mn concentration can be 10–100 times higher than in neurons. Recent results have identified nine molecularly distinct clusters of hippocampal astrocytes (De Ceglia, 2023), suggesting that the molecular heterogeneity of astrocytes could account for the observed differences in Mn accumulation. Further work is required to confirm the existence of a subtype of astrocyte with high Mn content, and to characterize the molecular mechanisms involved in this process.

While high levels of Mn internalized by astrocytes may limit exposure to neurons, this comes at the cost of astrocytes becoming the target of toxic effects. Overexposure to Mn has been shown to alter glutamatergic function mediated by astrocytes in the hippocampus of a young mouse model of Alzheimer’s disease (Spitznagel, 2023). Exposure to Mn in mice induced astrocyte dysfunction in the hippocampus, disrupting the glutamine–glutamate–GABA (Gln–Glu– GABA) metabolic cycle and reducing GABA synthesis (Bi, 2024). Mn exposure affects the activity of astrocytic glutamine synthetase and glutamate transporters, leading to impaired neurotransmitter cycling, excitotoxicity, and reactive astrocytes that can exacerbate neuroinflammation mediated by NF-κB and oxidative stress (Ke, 2019).

Overall, our results suggest that the primary mechanism of neuroprotection is Mn storage in the Golgi apparatus, involving both neurons and astrocytes. However, once the Golgi apparatus’s Mn storage capability is exceeded, it becomes a primary target of neurotoxicity. Mn can reach other critical intracellular regions, such as the postsynaptic compartments in neurons. The Golgi apparatus’s ability to detoxify Mn may be impaired by excessive exposure to Mn or by mutations in Mn efflux proteins such as SLC30A10 (Carmona et al., 2019).

Our study emphasizes the importance of early detection and intervention in neurological disorders related to exposure to Mn, particularly in occupational and environmental settings. The most effective way to avoid excessive exposure to Mn is to prevent exposure in the first place. Numerous studies have shown the neurotoxic effects of Mn in children and adults, and this has recently resulted in the WHO modifying its guideline on the recommended value of Mn in drinking water (WHO, 2022). While the previous guideline of 400 µg/L for Mn in drinking water was discontinued in 2012, the WHO recommended value was set at 80 µg/L in 2022, taking into account recent research on Mn neurotoxicology. This is a significant achievement in protecting the global population against Mn toxicity, as tens of millions of people currently drink water with a Mn concentration exceeding the WHO’s recommended value (Frisbie, 2012). Our study also emphasizes the need for further research into new therapeutic approaches for Mn detoxification that could enhance Mn efflux through the Golgi apparatus pathway. Such therapeutic interventions are urgently needed for pathologies associated with SLC30A10 and SLC39A14 mutations.

## Author Contributions

I.K. performed sample preparation, synchrotron experiments and data treatment. A.K. performed synchrotron experiments and data treatment. L.C.C.H. provided assistance during synchrotron experiments and data treatment. H.C. and M.S. developed the new ID21 setup, and provided assistance during synchrotron experiments and data analysis. S.R. and P.B. developed protocol, made measurements and performed data treatment of neurons viability assay. M.S. conceived the PSD95 fluorescent marker. N.P. and M.F.M designed cryo-FLM experiments and provided assistance. D.C. conceived the experiments. R.O. and A.C. conceived the experiments, performed synchrotron experiments, interpreted the results, and wrote the manuscript. All authors edited the final manuscript and approved the final version.

## Acknowledgment

The authors acknowledge the support of the French National Research Agency (ANR), under grant ANR-21-CE34-0011 (project SuperResMetToxSyn). We also acknowledge the European Synchrotron Radiation Facility (ESRF) for provision of synchrotron radiation facilities under proposal numbers LS3142 and LS3418. We would like to thank the Cell Biology Facility (PBC) of Interdisciplinary Institute of Neurosciences (IINS) at the University of Bordeaux, especially Nicolas Chevrier, Adrien Caralp and Natacha Retailleau, for general cell biology activity management. We acknowledge the Bordeaux Imaging Center (BIC), member of the national infrastructure France-BioImaging supported by the French National Research Agency (ANR-10-INBS-04).

## Notes

### Competing Interest Statement

The authors have declared no competing interest.

