## Supplementary material for "Synchrotron XRF imaging reveals manganese accumulation in the Golgi and post-synapses of neurons and enhanced uptake in astrocytes"

**Supplementary material 1**: **illustration of the correlative procedure using ICY and eC-CLEM.**

### Step 1: prepare merged images from cryo-FLM, freeze dry microscopy and SXRF maps

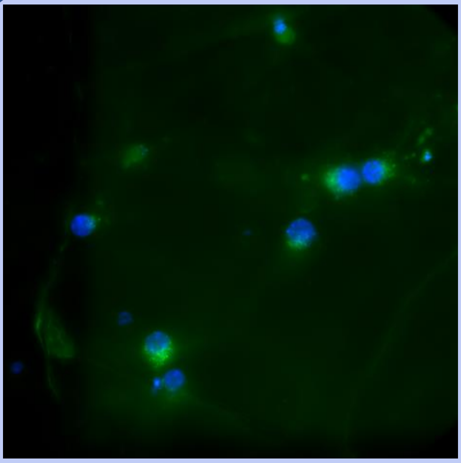

1. image composed of two channels, cryo-FLM of Golgi and nucleus.

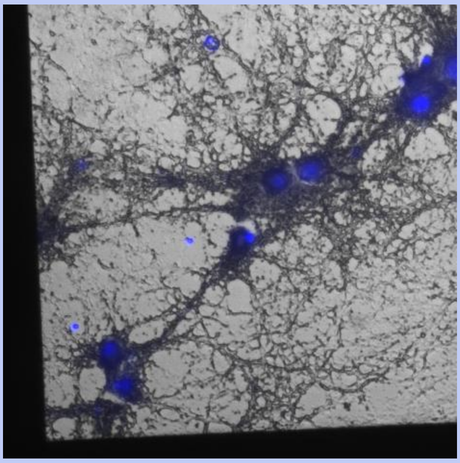

2. Image composed of two channels, nucleus and bright-field after freeze dry (usually GFP is not well preserved).

3. Compose a merged image containing all the SXRF maps of interest in different channels.

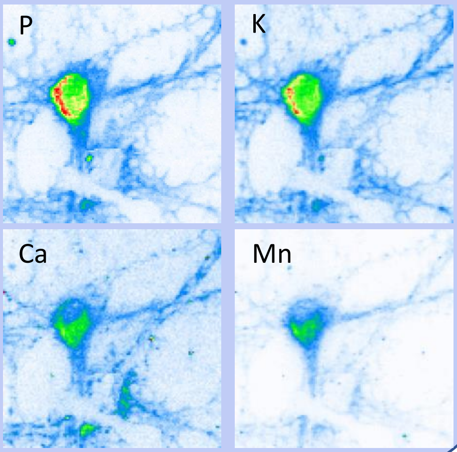

### Step 2: Align cryo-FLM and freeze dry multichannel images using eC-CLEM

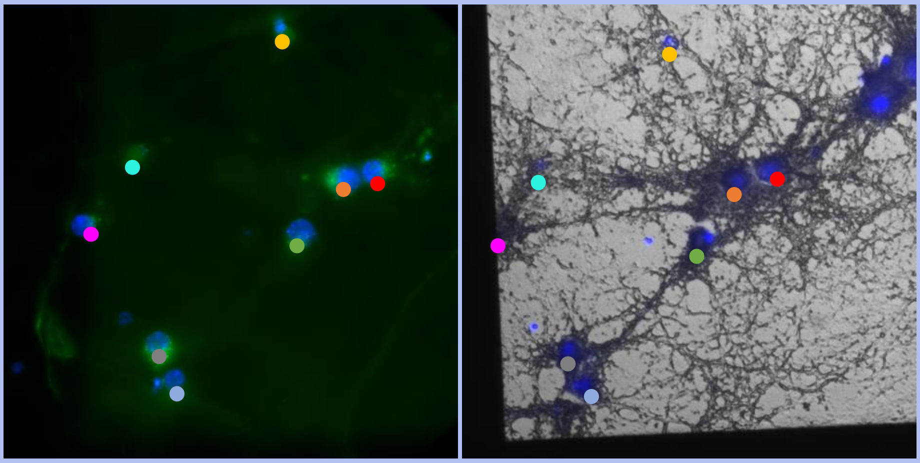

Alignment: the positions of the nuclei (cryo-FLM and freeze dry) are used to affine sample registration.

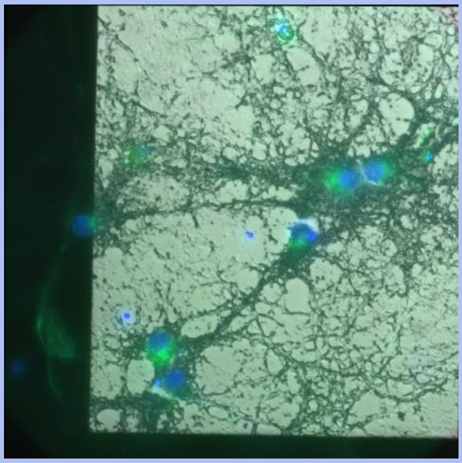

Final aligned image with 4 channels: 2 from cryo-FLM and 2 from freeze dry.

### Step 3: Align multichannel SXRF image and the final image from step 2

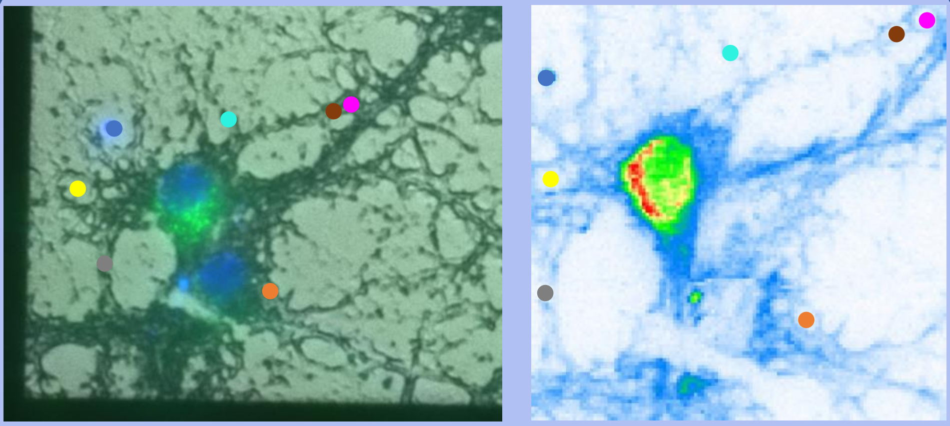

To affine sample registration and precise alignment, use the shape of the dendrites, visualized with the bright-field channel and P or K channels.

### Step 4: Visualize the region of interest

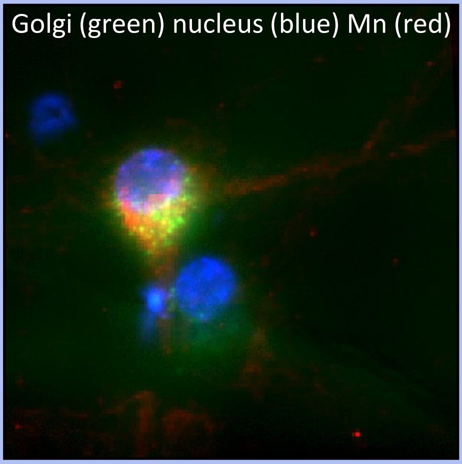

The final image is composed of different channels containing cryo-FLM, freeze-dried and SXRF images. Select the channels of interest to visualize the correlation between SXRF and cryo-FLM.

**Supplementary material 2**: quantitative data expressed in $ng/mm^2$.

| $ng/mm^2$ | P | | | S | | | K | | | Ca | | | Mn | | |
|---|---|---|---|---|---|---|---|---|---|---|---|---|---|---|---|
| data | Neurons | Mixed Neurons | Astrocytes | Neurons | Mixed Neurons | Astrocytes | Neurons | Mixed Neurons | Astrocytes | Neurons | Mixed Neurons | Astrocytes | Neurons | Mixed Neurons | Astrocytes |
| | 11.6 | 29.7 | 5.16 | 6.13 | 8.42 | 2.3 | 1.37 | 13.84 | 4 | 0.66 | 2.95 | 0.38 | 0.29 | 1.28 | 1.23 |
| | 27.39 | 17.56 | 18.87 | 7.84 | 5.1 | 5.27 | 2.89 | 8.61 | 9.57 | 0.57 | 0.7 | 0.54 | 0.66 | 0.42 | 0.36 |
| | 6.47 | 14.82 | 24.39 | 2.27 | 5.05 | 8.59 | 0.86 | 7.25 | 16.94 | 0.18 | 0.2 | 0.39 | 0.23 | 0.16 | 0.23 |
| | 25.71 | 4.47 | 27.13 | 7.39 | 1.52 | 8.2 | 1.84 | 0.67 | 16.11 | 0.66 | 0.28 | 0.42 | 0.87 | 0.13 | 0.55 |
| | 21.84 | 14.06 | 12.82 | 6.25 | 4.29 | 4.17 | 1.9 | 7.78 | 8.1 | 0.65 | 0.49 | 0.2 | 0.97 | 0.36 | 0.39 |
| | 24.14 | 33.75 | 15.54 | 7 | 9.82 | 5.51 | 2.05 | 16.42 | 12.31 | 0.73 | 0.42 | 0.34 | 0.75 | 0.47 | 0.66 |
| | 33.7 | 27.93 | 7.19 | 9.4 | 8.1 | 2.57 | 3.8 | 13.18 | 5.79 | 0.73 | 0.44 | 0.14 | 1.02 | 0.44 | 0.26 |
| | 27.26 | 25.93 | 8.53 | 5.36 | 7.69 | 3.04 | 4.07 | 13.37 | 7.97 | 3.93 | 0.58 | 0.17 | 13.06 | 0.4 | 0.5 |
| | 4.79 | 17.28 | 4.97 | 1.65 | 4.72 | 1.83 | 0.56 | 8.02 | 4.33 | 0.15 | 0.89 | 0.21 | 0.16 | 0.44 | 0.23 |
| | 9.6 | 21.22 | 11.12 | 3.11 | 6.29 | 4.09 | 1.1 | 7.71 | 7.69 | 0.26 | 0.74 | 0.34 | 0.28 | 0.36 | 0.41 |
| | 18.5 | 19.96 | 10.36 | 4.86 | 6.01 | 4.02 | 1.29 | 8.62 | 8.73 | 0.31 | 0.42 | 0.3 | 0.45 | 0.24 | 0.44 |
| | 21.48 | 21.22 | 24.27 | 5.25 | 5.83 | 6.54 | 2.95 | 8.54 | 15.54 | 0.46 | 0.94 | 1 | 0.83 | 0.47 | 9.19 |
| | 31.17 | 36.46 | 8.81 | 9.53 | 11.82 | 2.42 | 2.83 | 17.61 | 4.11 | 0.73 | 0.73 | 0.71 | 0.79 | 0.4 | 3.71 |
| | 40.72 | 41.2 | 8.74 | 13.07 | 17.19 | 2.77 | 3.31 | 17.9 | 6.31 | 0.9 | 1.57 | 0.18 | 1.01 | 0.45 | 0.5 |
| | 44.36 | 55.27 | 24.88 | 14.11 | 19.19 | 7.05 | 3.02 | 25.03 | 4.27 | 0.85 | 1.76 | 0.88 | 0.99 | 0.7 | 0.52 |
| | 27.82 | | 26.47 | 8.73 | | 6.62 | 2.52 | | 3.62 | 0.55 | | 1.64 | 0.68 | | 0.09 |
| | 16.65 | | 4.77 | 5.6 | | 1.88 | 1.51 | | 15.23 | 0.44 | | 0.21 | 0.28 | | 5.05 |
| | 8.07 | | 3.95 | 2.73 | | 1.61 | 0.92 | | 9.99 | 0.23 | | 0.09 | 0.16 | | 15.95 |
| | 27.85 | | 24.96 | 8.81 | | 10.59 | 2.08 | | 17.17 | 0.63 | | 0.37 | 0.67 | | 0.25 |
| | 19.9 | | | 6.81 | | | 1.79 | | | 0.55 | | | 0.5 | | |
| | 51.16 | | | 16.89 | | | 5.89 | | | 1.05 | | | 1.31 | | |
| | 29.99 | | | 9.52 | | | 2.04 | | | 0.62 | | | 0.7 | | |
| | 56.22 | | | 22.51 | | | 9.02 | | | 0.83 | | | 1.12 | | |
| Number of values | 23 | 15 | 19 | 23 | 15 | 19 | 23 | 15 | 19 | 23 | 15 | 19 | 23 | 15 | 19 |
| Minimum | 4.8 | 4.5 | 4.0 | 1.7 | 1.5 | 1.6 | 0.56 | 0.67 | 3.6 | 0.15 | 0.20 | 0.090 | 0.16 | 0.13 | 0.090 |
| 25% Percentile | 17 | 17 | 7.2 | 5.3 | 5.1 | 2.4 | 1.4 | 7.8 | 4.3 | 0.44 | 0.42 | 0.20 | 0.29 | 0.36 | 0.26 |
| Median | 26 | 21 | 11 | 7.0 | 6.3 | 4.1 | 2.1 | 8.6 | 8.1 | 0.63 | 0.70 | 0.34 | 0.70 | 0.42 | 0.50 |
| 75% Percentile | 31 | 34 | 24 | 9.5 | 9.8 | 6.6 | 3.0 | 16 | 15 | 0.73 | 0.94 | 0.54 | 0.99 | 0.47 | 1.2 |
| Maximum | 56 | 55 | 27 | 23 | 19 | 11 | 9.0 | 25 | 17 | 3.9 | 3.0 | 1.6 | 13 | 1.3 | 16 |
| Range | 51 | 51 | 23 | 21 | 18 | 9.0 | 8.5 | 24 | 14 | 3.8 | 2.8 | 1.6 | 13 | 1.2 | 16 |
| 95% CI of median | | | | | | | | | | | | | | | |
| Actual confidence level | 97% | 96% | 98% | 97% | 96% | 98% | 97% | 96% | 98% | 97% | 96% | 98% | 97% | 96% | 98% |
| Lower confidence limit | 19 | 17 | 7.2 | 5.4 | 5.1 | 2.4 | 1.5 | 7.8 | 4.3 | 0.46 | 0.42 | 0.20 | 0.45 | 0.36 | 0.26 |
| Upper confidence limit | 30 | 34 | 24 | 9.4 | 9.8 | 6.6 | 3.0 | 16 | 15 | 0.73 | 0.94 | 0.54 | 0.97 | 0.47 | 1.2 |

**Supplementary material 3**: **elemental concentrations expressed in mM.**

$$C_{vol}\ [mM] = \frac{C_{surf}\ [ng/mm^2]}{cell\ thickness\ [mm] \cdot MW\ [g/mol]}$$

The [cell thickness] was measured in living conditions, hydrated cells, and MW is the molecular weight of the element of interest.

| mM | P | | | S | | | K | | | Ca | | | Mn | | |
|---|---|---|---|---|---|---|---|---|---|---|---|---|---|---|---|
| data | Neurons | Mixed Neurons | Astrocytes | Neurons | Mixed Neurons | Astrocytes | Neurons | Mixed Neurons | Astrocytes | Neurons | Mixed Neurons | Astrocytes | Neurons | Mixed Neurons | Astrocytes |
| | 24.16 | 61.86 | 28.72 | 12.33 | 16.94 | 12.37 | 2.26 | 22.84 | 17.64 | 1.06 | 4.75 | 1.63 | 0.34 | 1.50 | 3.86 |
| | 57.05 | 36.58 | 105.04 | 15.77 | 10.26 | 28.33 | 4.77 | 14.21 | 42.20 | 0.92 | 1.13 | 2.32 | 0.78 | 0.49 | 1.13 |
| | 13.48 | 30.87 | 135.77 | 4.57 | 10.16 | 46.18 | 1.42 | 11.96 | 74.70 | 0.29 | 0.32 | 1.68 | 0.27 | 0.19 | 0.72 |
| | 53.55 | 9.31 | 151.02 | 14.87 | 3.06 | 44.09 | 3.04 | 1.11 | 71.04 | 1.06 | 0.45 | 1.81 | 1.02 | 0.15 | 1.73 |
| | 45.49 | 29.29 | 71.36 | 12.57 | 8.63 | 22.42 | 3.14 | 12.84 | 35.72 | 1.05 | 0.79 | 0.86 | 1.14 | 0.42 | 1.22 |
| | 50.28 | 70.30 | 86.50 | 14.08 | 19.76 | 29.63 | 3.38 | 27.09 | 54.28 | 1.18 | 0.68 | 1.46 | 0.88 | 0.55 | 2.07 |
| | 70.20 | 58.18 | 40.02 | 18.91 | 16.30 | 13.82 | 6.27 | 21.75 | 25.53 | 1.18 | 0.71 | 0.60 | 1.20 | 0.52 | 0.82 |
| | 56.78 | 54.01 | 47.48 | 10.78 | 15.47 | 16.34 | 6.72 | 22.06 | 35.15 | 6.33 | 0.93 | 0.73 | 15.34 | 0.47 | 1.57 |
| | 9.98 | 35.99 | 27.67 | 3.32 | 9.50 | 9.84 | 0.92 | 13.23 | 19.09 | 0.24 | 1.43 | 0.90 | 0.19 | 0.52 | 0.72 |
| | 20.00 | 44.20 | 61.90 | 6.26 | 12.65 | 21.99 | 1.82 | 12.72 | 33.91 | 0.42 | 1.19 | 1.46 | 0.33 | 0.42 | 1.29 |
| | 38.54 | 41.58 | 57.67 | 9.78 | 12.09 | 21.61 | 2.13 | 14.22 | 38.50 | 0.50 | 0.68 | 1.29 | 0.53 | 0.28 | 1.38 |
| | 44.74 | 44.20 | 135.10 | 10.56 | 11.73 | 35.16 | 4.87 | 14.09 | 68.53 | 0.74 | 1.51 | 4.30 | 0.97 | 0.55 | 28.84 |
| | 64.93 | 75.95 | 49.04 | 19.17 | 23.78 | 13.01 | 4.67 | 29.06 | 18.12 | 1.18 | 1.18 | 3.05 | 0.93 | 0.47 | 11.64 |
| | 84.82 | 85.82 | 48.65 | 26.30 | 34.58 | 14.89 | 5.46 | 29.54 | 27.83 | 1.45 | 2.53 | 0.77 | 1.19 | 0.53 | 1.57 |
| | 92.40 | 115.13 | 138.50 | 28.39 | 38.61 | 37.90 | 4.98 | 41.30 | 18.83 | 1.37 | 2.83 | 3.79 | 1.16 | 0.82 | 1.63 |
| | 57.95 | | 147.35 | 17.56 | | 35.59 | 4.16 | | 15.96 | 0.89 | | 7.06 | 0.80 | | 0.28 |
| | 34.68 | | 26.55 | 11.27 | | 10.11 | 2.49 | | 67.16 | 0.71 | | 0.90 | 0.33 | | 15.85 |
| | 16.81 | | 21.99 | 5.49 | | 8.66 | 1.52 | | 44.05 | 0.37 | | 0.39 | 0.19 | | 50.06 |
| | 58.01 | | 138.94 | 17.72 | | 56.94 | 3.43 | | 75.72 | 1.01 | | 1.59 | 0.79 | | 0.78 |
| | 41.45 | | | 13.70 | | | 2.95 | | | 0.89 | | | 0.59 | | |
| | 106.57 | | | 33.98 | | | 9.72 | | | 1.69 | | | 1.54 | | |
| | 62.47 | | | 19.15 | | | 3.37 | | | 1.00 | | | 0.82 | | |
| | 117.11 | | | 45.29 | | | 14.88 | | | 1.34 | | | 1.32 | | |
| Number of values | 23 | 15 | 19 | 23 | 15 | 19 | 23 | 15 | 19 | 23 | 15 | 19 | 23 | 15 | 19 |
| Minimum | 10 | 9.3 | 22 | 3.3 | 3.1 | 8.7 | 0.92 | 1.1 | 16 | 0.24 | 0.32 | 0.39 | 0.19 | 0.15 | 0.28 |
| 25% Percentile | 35 | 36 | 40 | 11 | 10 | 13 | 2.3 | 13 | 19 | 0.71 | 0.68 | 0.86 | 0.34 | 0.42 | 0.82 |
| Median | 54 | 44 | 62 | 14 | 13 | 22 | 3.4 | 14 | 36 | 1.0 | 1.1 | 1.5 | 0.82 | 0.49 | 1.6 |
| 75% Percentile | 65 | 70 | 136 | 19 | 20 | 36 | 5.0 | 27 | 67 | 1.2 | 1.5 | 2.3 | 1.2 | 0.55 | 3.9 |
| Maximum | 117 | 115 | 151 | 45 | 39 | 57 | 15 | 41 | 76 | 6.3 | 4.7 | 7.1 | 15 | 1.5 | 50 |
| Range | 107 | 106 | 129 | 42 | 36 | 48 | 14 | 40 | 60 | 6.1 | 4.4 | 6.7 | 15 | 1.4 | 50 |
| 95% CI of median | | | | | | | | | | | | | | | |
| Actual confidence level | 97% | 96% | 98% | 97% | 96% | 98% | 97% | 96% | 98% | 97% | 96% | 98% | 97% | 96% | 98% |
| Lower confidence limit | 39 | 36 | 40 | 11 | 10 | 13 | 2.5 | 13 | 19 | 0.74 | 0.68 | 0.86 | 0.53 | 0.42 | 0.82 |
| Upper confidence limit | 62 | 70 | 136 | 19 | 20 | 36 | 4.9 | 27 | 67 | 1.2 | 1.5 | 2.3 | 1.1 | 0.55 | 3.9 |

**Supplementary material 4: measured thickness of neurons and astrocytes in living conditions.**

| | neurons thickness (µm) | astrocytes thickness (µm) |
|---|---|---|
| | 17.0 | 5.4 |
| | 11.0 | 5.7 |
| | 16.0 | 6.6 |
| | 16.0 | 6.0 |
| | 16.0 | 5.8 |
| | 16.0 | 7.6 |
| | 16.0 | 5.2 |
| | 16.0 | 6.0 |
| | 16.0 | 5.5 |
| | 13.0 | 5.8 |
| | 19.0 | 5.4 |
| | 14.0 | 5.2 |
| | 16.7 | 6.5 |
| | 9.0 | 7.0 |
| | 15.6 | 5.4 |
| | 13.4 | 4.8 |
| | 16.0 | 5.0 |
| | 15.9 | 5.8 |
| | 16.7 | 6.8 |
| | 17.6 | 4.2 |
| | 18.3 | 6.0 |
| **Mean (µm)** | 15.5 | 5.8 |
| Std Dev | 2.3 | 0.8 |

**Supplementary material 5 (P): statistics from supplementary material 3, expressed in mM for P element.**

| P | | | | | |
|---|---|---|---|---|---|
| Table Analyzed | normalized (mM) | Table Analyzed | normalized (mM) | Table Analyzed | normalized (mM) |
| Column C | Mixed Neurons | Column D | Astrocytes | Column D | Astrocytes |
| vs. | vs. | vs. | vs. | vs. | vs. |
| Column B | Neurons | Column B | Neurons | Column C | Mixed Neurons |
| Mann Whitney test | | Mann Whitney test | | Mann Whitney test | |
| P value | 0.9472 | P value | 0.1009 | P value | 0.1647 |
| Exact or approximate P value? | Exact | Exact or approximate P value? | Exact | Exact or approximate P value? | Exact |
| P value summary | ns | P value summary | ns | P value summary | ns |
| Significantly different (P < 0.05)? | No | Significantly different (P < 0.05)? | No | Significantly different (P < 0.05)? | No |
| One- or two-tailed P value? | Two-tailed | One- or two-tailed P value? | Two-tailed | One- or two-tailed P value? | Two-tailed |
| Sum of ranks in column B,C | 451 , 290 | Sum of ranks in column B,D | 429 , 474 | Sum of ranks in column C,D | 222 , 373 |
| Mann-Whitney U | 170 | Mann-Whitney U | 153 | Mann-Whitney U | 102 |
| Difference between medians | | Difference between medians | | Difference between medians | |
| Median of column B | 53.55, n=23 | Median of column B | 53.55, n=23 | Median of column C | 44.20, n=15 |
| Median of column C | 44.20, n=15 | Median of column D | 61.90, n=19 | Median of column D | 61.90, n=19 |
| Difference: Actual | -9.353 | Difference: Actual | 8.347 | Difference: Actual | 17.70 |
| Difference: Hodges-Lehmann | -0.6086 | Difference: Hodges-Lehmann | 19.13 | Difference: Hodges-Lehmann | 18.20 |
| 95.22% CI of difference | -17.56 to 18.89 | 95.21% CI of difference | -4.799 to 51.82 | 95.29% CI of difference | -7.299 to 61.53 |
| Exact or approximate CI? | Exact | Exact or approximate CI? | Exact | Exact or approximate CI? | Exact |

**Supplementary material 5 (S): statistics from supplementary material 3, expressed in mM for S element.**

| S | | | | | |
|---|---|---|---|---|---|
| Table Analyzed | normalized (mM) | Table Analyzed | normalized (mM) | Table Analyzed | normalized (mM) |
| Column G | Mixed Neurons | Column H | Astrocytes | Column H | Astrocytes |
| vs. | vs. | vs. | vs. | vs. | vs. |
| Column F | Neurons | Column F | Neurons | Column G | Mixed Neurons |
| Mann Whitney test | | Mann Whitney test | | Mann Whitney test | |
| P value | 0.8828 | P value | 0.0327 | P value | 0.0557 |
| Exact or approximate P value? | Exact | Exact or approximate P value? | Exact | Exact or approximate P value? | Exact |
| P value summary | ns | P value summary | * | P value summary | ns |
| Significantly different (P < 0.05)? | No | Significantly different (P < 0.05)? | Yes | Significantly different (P < 0.05)? | No |
| One- or two-tailed P value? | Two-tailed | One- or two-tailed P value? | Two-tailed | One- or two-tailed P value? | Two-tailed |
| Sum of ranks in column F,G | 454 , 287 | Sum of ranks in column F,H | 410 , 493 | Sum of ranks in column G,H | 207 , 388 |
| Mann-Whitney U | 167 | Mann-Whitney U | 134 | Mann-Whitney U | 87 |
| Difference between medians | | Difference between medians | | Difference between medians | |
| Median of column F | 14.08, n=23 | Median of column F | 14.08, n=23 | Median of column G | 12.65, n=15 |
| Median of column G | 12.65, n=15 | Median of column H | 21.99, n=19 | Median of column H | 21.99, n=19 |
| Difference: Actual | -1.428 | Difference: Actual | 7.907 | Difference: Actual | 9.335 |
| Difference: Hodges-Lehmann | -0.4024 | Difference: Hodges-Lehmann | 7.531 | Difference: Hodges-Lehmann | 6.262 |
| 95.22% CI of difference | -5.372 to 5.472 | 95.21% CI of difference | 0.5717 to 16.50 | 95.29% CI of difference | -0.1526 to 18.17 |
| Exact or approximate CI? | Exact | Exact or approximate CI? | Exact | Exact or approximate CI? | Exact |

**Supplementary material 5 (K): statistics from supplementary material 3, expressed in mM for K element.**

| K | | | | | |
|---|---|---|---|---|---|
| Table Analyzed | normalized (mM) | Table Analyzed | normalized (mM) | Table Analyzed | normalized (mM) |
| Column K | Mixed Neurons | Column L | Astrocytes | Column L | Astrocytes |
| vs. | vs. | vs. | vs. | vs. | vs. |
| Column J | Neurons | Column J | Neurons | Column K | Mixed Neurons |
| Mann Whitney test | | Mann Whitney test | | Mann Whitney test | |
| P value | <0.0001 | P value | <0.0001 | P value | 0.0005 |
| Exact or approximate P value? | Exact | Exact or approximate P value? | Exact | Exact or approximate P value? | Exact |
| P value summary | **** | P value summary | **** | P value summary | *** |
| Significantly different (P < 0.05)? | Yes | Significantly different (P < 0.05)? | Yes | Significantly different (P < 0.05)? | Yes |
| One- or two-tailed P value? | Two-tailed | One- or two-tailed P value? | Two-tailed | One- or two-tailed P value? | Two-tailed |
| Sum of ranks in column J,K | 305 , 436 | Sum of ranks in column J,L | 276 , 627 | Sum of ranks in column K,L | 166 , 429 |
| Mann-Whitney U | 29 | Mann-Whitney U | 0 | Mann-Whitney U | 46 |
| Difference between medians | | Difference between medians | | Difference between medians | |
| Median of column J | 3.383, n=23 | Median of column J | 3.383, n=23 | Median of column K | 14.22, n=15 |
| Median of column K | 14.22, n=15 | Median of column L | 35.72, n=19 | Median of column L | 35.72, n=19 |
| Difference: Actual | 10.84 | Difference: Actual | 32.34 | Difference: Actual | 21.50 |
| Difference: Hodges-Lehmann | 12.03 | Difference: Hodges-Lehmann | 32.49 | Difference: Hodges-Lehmann | 19.70 |
| 95.22% CI of difference | 9.587 to 19.70 | 95.21% CI of difference | 22.96 to 42.24 | 95.29% CI of difference | 5.992 to 32.81 |
| Exact or approximate CI? | Exact | Exact or approximate CI? | Exact | Exact or approximate CI? | Exact |

**Supplementary material 5 (Ca): statistics from supplementary material 3, expressed in mM for Ca element.**

| Ca | | | | | |
|---|---|---|---|---|---|
| Table Analyzed | normalized (mM) | Table Analyzed | normalized (mM) | Table Analyzed | normalized (mM) |
| Column O | Mixed Neurons | Column P | Astrocytes | Column P | Astrocytes |
| vs. | vs. | vs. | vs. | vs. | vs. |
| Column N | Neurons | Column N | Neurons | Column O | Mixed Neurons |
| Mann Whitney test | | Mann Whitney test | | Mann Whitney test | |
| P value | 0.5107 | P value | 0.0343 | P value | 0.1991 |
| Exact or approximate P value? | Exact | Exact or approximate P value? | Exact | Exact or approximate P value? | Exact |
| P value summary | ns | P value summary | * | P value summary | ns |
| Significantly different (P < 0.05)? | No | Significantly different (P < 0.05)? | Yes | Significantly different (P < 0.05)? | No |
| One- or two-tailed P value? | Two-tailed | One- or two-tailed P value? | Two-tailed | One- or two-tailed P value? | Two-tailed |
| Sum of ranks in column N,O | 426 , 315 | Sum of ranks in column N,P | 411 , 492 | Sum of ranks in column O,P | 225 , 370 |
| Mann-Whitney U | 150 | Mann-Whitney U | 135 | Mann-Whitney U | 105 |
| Difference between medians | | Difference between medians | | Difference between medians | |
| Median of column N | 1.014, n=23 | Median of column N | 1.014, n=23 | Median of column O | 1.127, n=15 |
| Median of column O | 1.127, n=15 | Median of column P | 1.463, n=19 | Median of column P | 1.463, n=19 |
| Difference: Actual | 0.1127 | Difference: Actual | 0.4485 | Difference: Actual | 0.3358 |
| Difference: Hodges-Lehmann | 0.1127 | Difference: Hodges-Lehmann | 0.4707 | Difference: Hodges-Lehmann | 0.2875 |
| 95.22% CI of difference | -0.2415 to 0.5151 | 95.21% CI of difference | 0.01804 to 0.9636 | 95.29% CI of difference | -0.2664 to 0.9586 |
| Exact or approximate CI? | Exact | Exact or approximate CI? | Exact | Exact or approximate CI? | Exact |

**Supplementary material 5 (Mn): statistics from supplementary material 3, expressed in mM for Mn element.**

| Mn | | | | | |
|---|---|---|---|---|---|
| Table Analyzed | normalized (mM) | Table Analyzed | normalized (mM) | Table Analyzed | normalized (mM) |
| Column S | Mixed Neurons | Column T | Astrocytes | Column T | Astrocytes |
| vs. | vs. | vs. | vs. | vs. | vs. |
| Column R | Neurons | Column R | Neurons | Column S | Mixed Neurons |
| Mann Whitney test | | Mann Whitney test | | Mann Whitney test | |
| P value | 0.0273 | P value | 0.0018 | P value | <0.0001 |
| Exact or approximate P value? | Exact | Exact or approximate P value? | Exact | Exact or approximate P value? | Exact |
| P value summary | * | P value summary | ** | P value summary | **** |
| Significantly different (P < 0.05)? | Yes | Significantly different (P < 0.05)? | Yes | Significantly different (P < 0.05)? | Yes |
| One- or two-tailed P value? | Two-tailed | One- or two-tailed P value? | Two-tailed | One- or two-tailed P value? | Two-tailed |
| Sum of ranks in column R,S | 522 , 219 | Sum of ranks in column R,T | 374 , 529 | Sum of ranks in column S,T | 144 , 451 |
| Mann-Whitney U | 99 | Mann-Whitney U | 98 | Mann-Whitney U | 24 |
| Difference between medians | | Difference between medians | | Difference between medians | |
| Median of column R | 0.8220, n=23 | Median of column R | 0.8220, n=23 | Median of column S | 0.4932, n=15 |
| Median of column S | 0.4932, n=15 | Median of column T | 1.569, n=19 | Median of column T | 1.569, n=19 |
| Difference: Actual | -0.3288 | Difference: Actual | 0.7471 | Difference: Actual | 1.076 |
| Difference: Hodges-Lehmann | -0.3406 | Difference: Hodges-Lehmann | 0.6572 | Difference: Hodges-Lehmann | 1.041 |
| 95.22% CI of difference | -0.6107 to -0.03523 | 95.21% CI of difference | 0.3078 to 1.273 | 95.29% CI of difference | 0.5779 to 1.555 |
| Exact or approximate CI? | Exact | Exact or approximate CI? | Exact | Exact or approximate CI? | Exact |

**Supplementary material 6: SXRF imaging in neurons and astrocytes not exposed to manganese.** a) Elemental distribution of P, S, K, Ca and Mn obtained by SXRF in single neuron, scan size 65 µm x 65 µm, step 0.5 µm/pixel, scan time 100 ms. b) Elemental distribution of P, S, K, Ca and Mn obtained by SXRF in single astrocytes, scan size 70 µm x 70 µm, step 0.5 µm/pixel, scan time 100 ms. Scale bars: 5 µm.

a) SXRF maps in single neuron

P S K Ca Mn

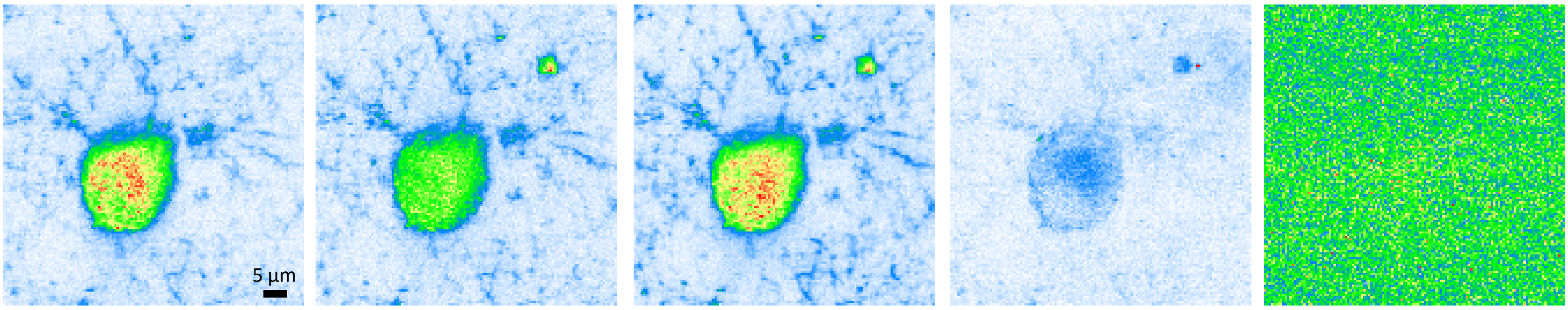

b) SXRF maps in a single astrocyte

P S K Ca Mn

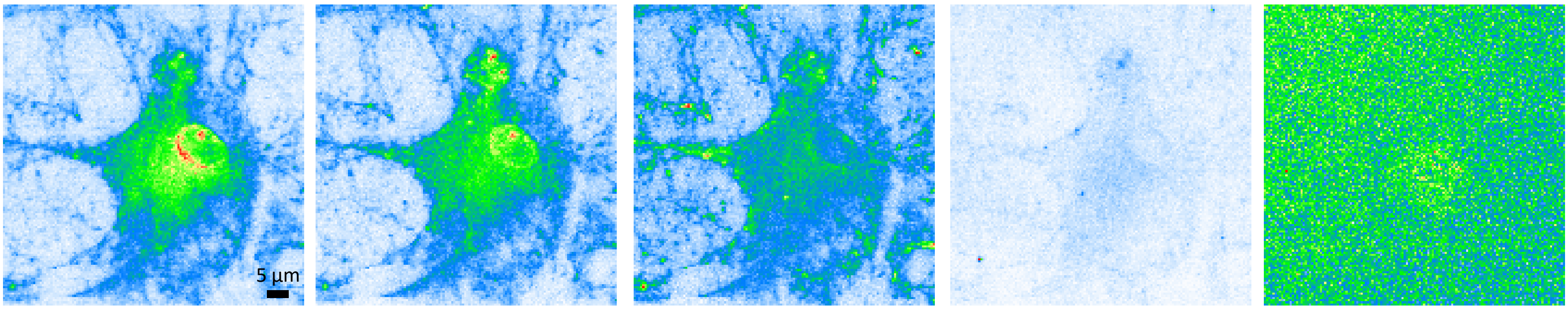

**Supplementary material 7: Manganese cytotoxicity assays.** Viability assay was performed in cultured primary neurons using the ReadyProbes™ Cell Viability Imaging Kit, Blue/Green. Manganese was added to the culture medium for 24h at different concentrations: 0-1-10-50-100-250-500-1000 µM. Normalized neuron viability, $IC_{10}$ (inhibitory concentration 10%) and $IC_{50}$ are shown. The plot presents three dose-response curves calculated by different models (Log-Logistic (LL4), Weibull Model 1 (W14), Weibull Model 2 (W24)) and the calculated weighted mean and the associated uncertainty (green curve). The viability was normalized to the control. The $IC_{10}$ (249 µM) and $IC_{50}$ (502 µM) values were calculated from the mean curve. n, is the total number of cells analyzed (131,472).

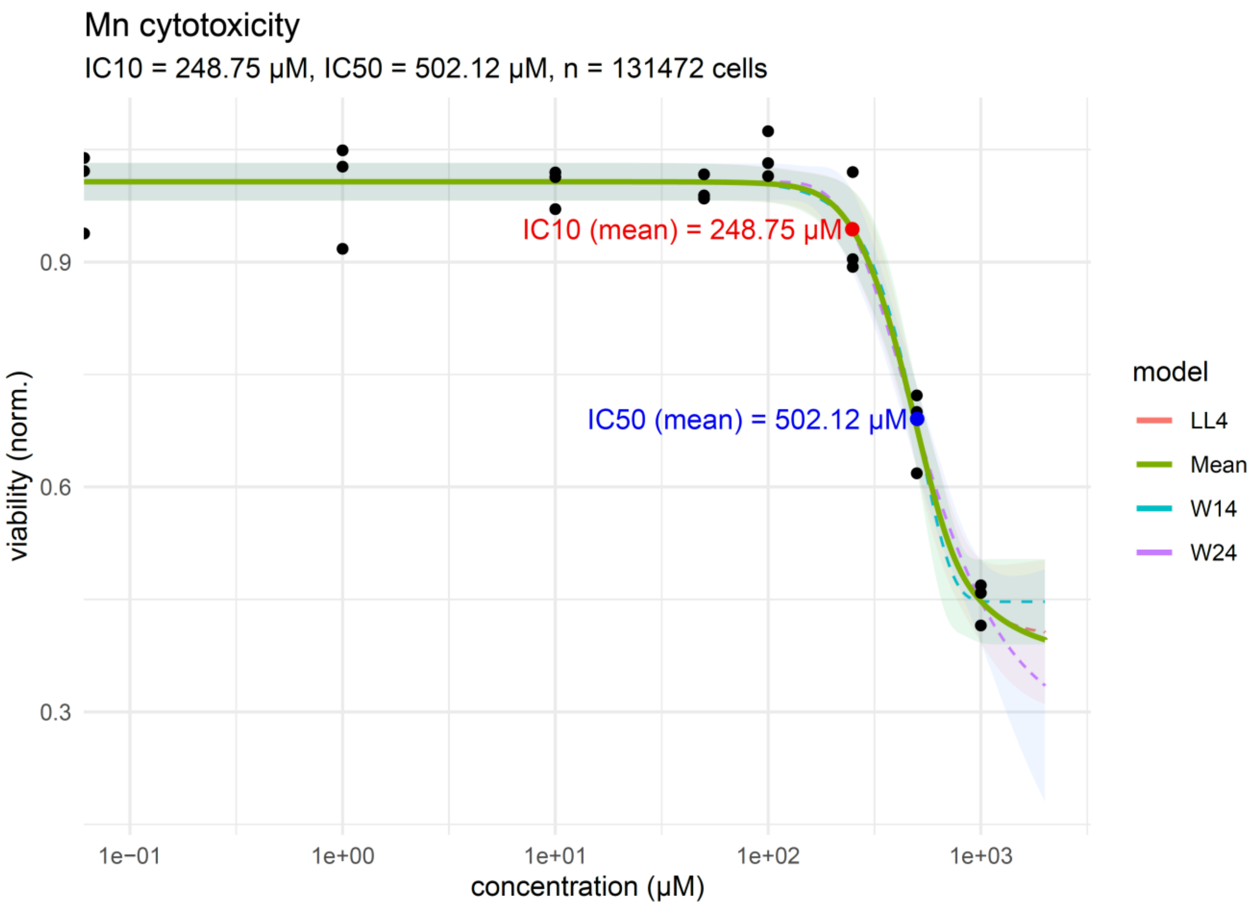

**Supplementary material 8: SXRF imaging in astrocytes exposed to manganese.**

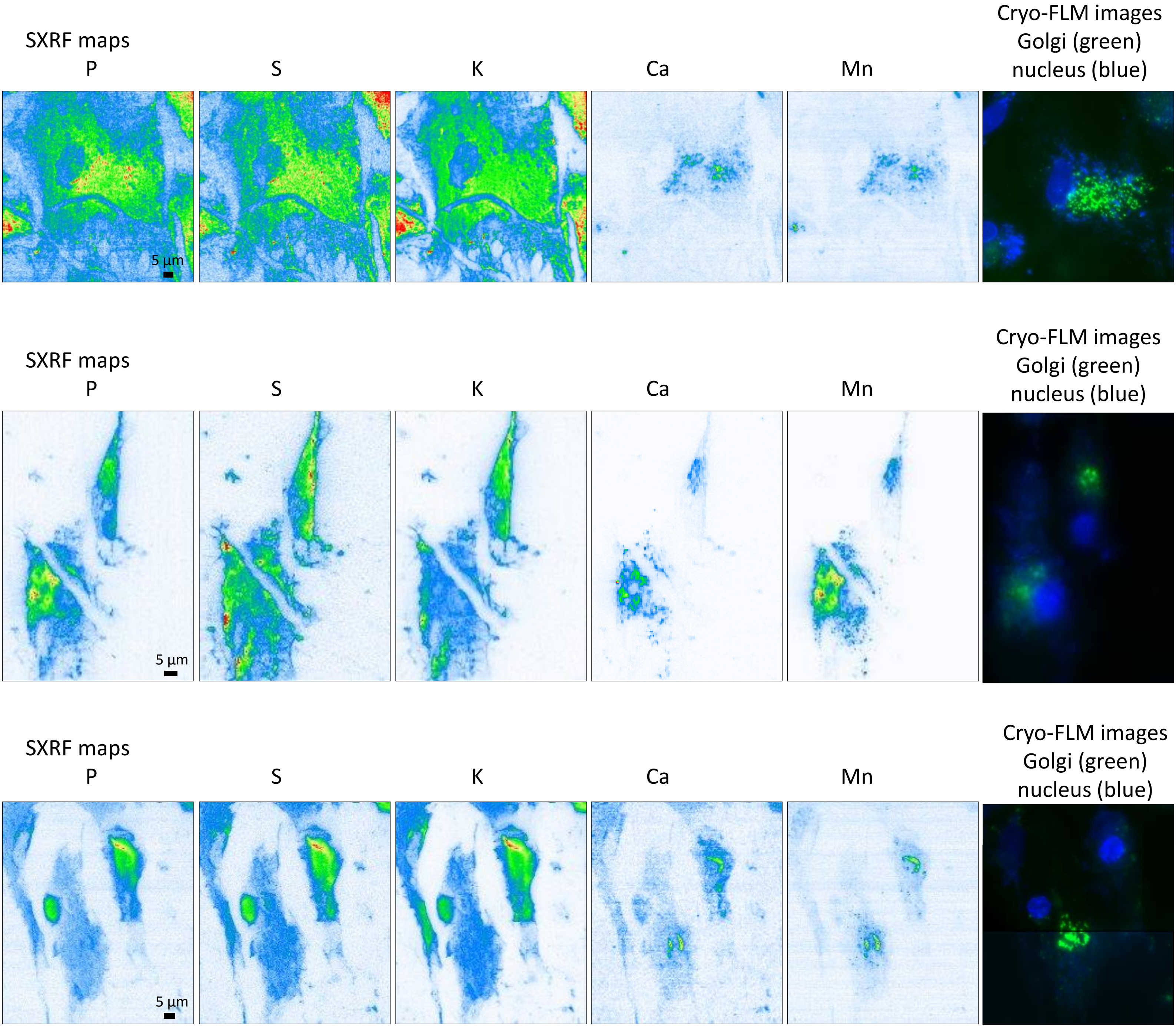

**Supplementary material 9**: **SXRF imaging in neurons exposed to manganese in the presence of astrocytes.**

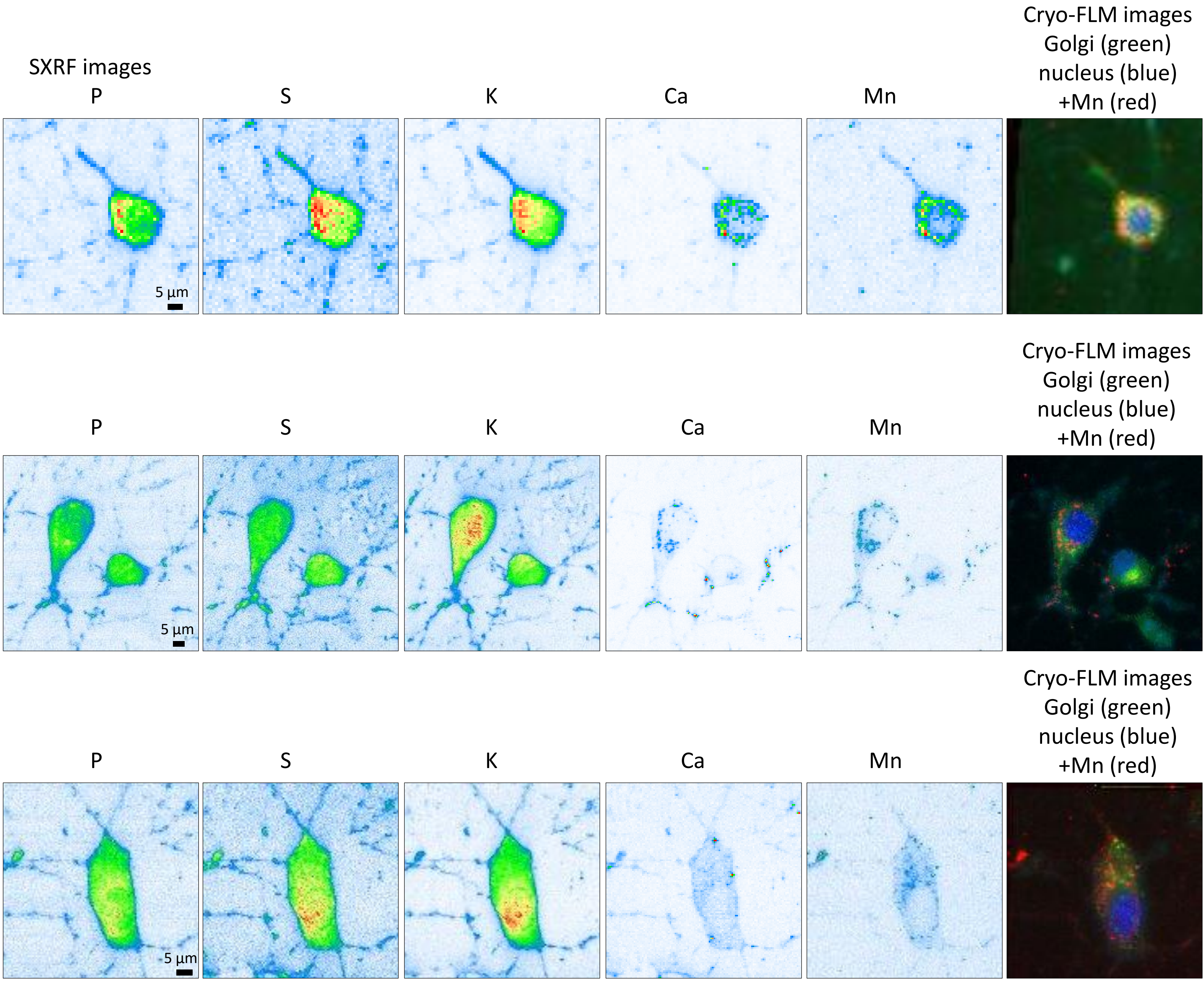

**Supplementary material 10: SXRF imaging in neurons exposed to manganese in the absence of astrocytes.**

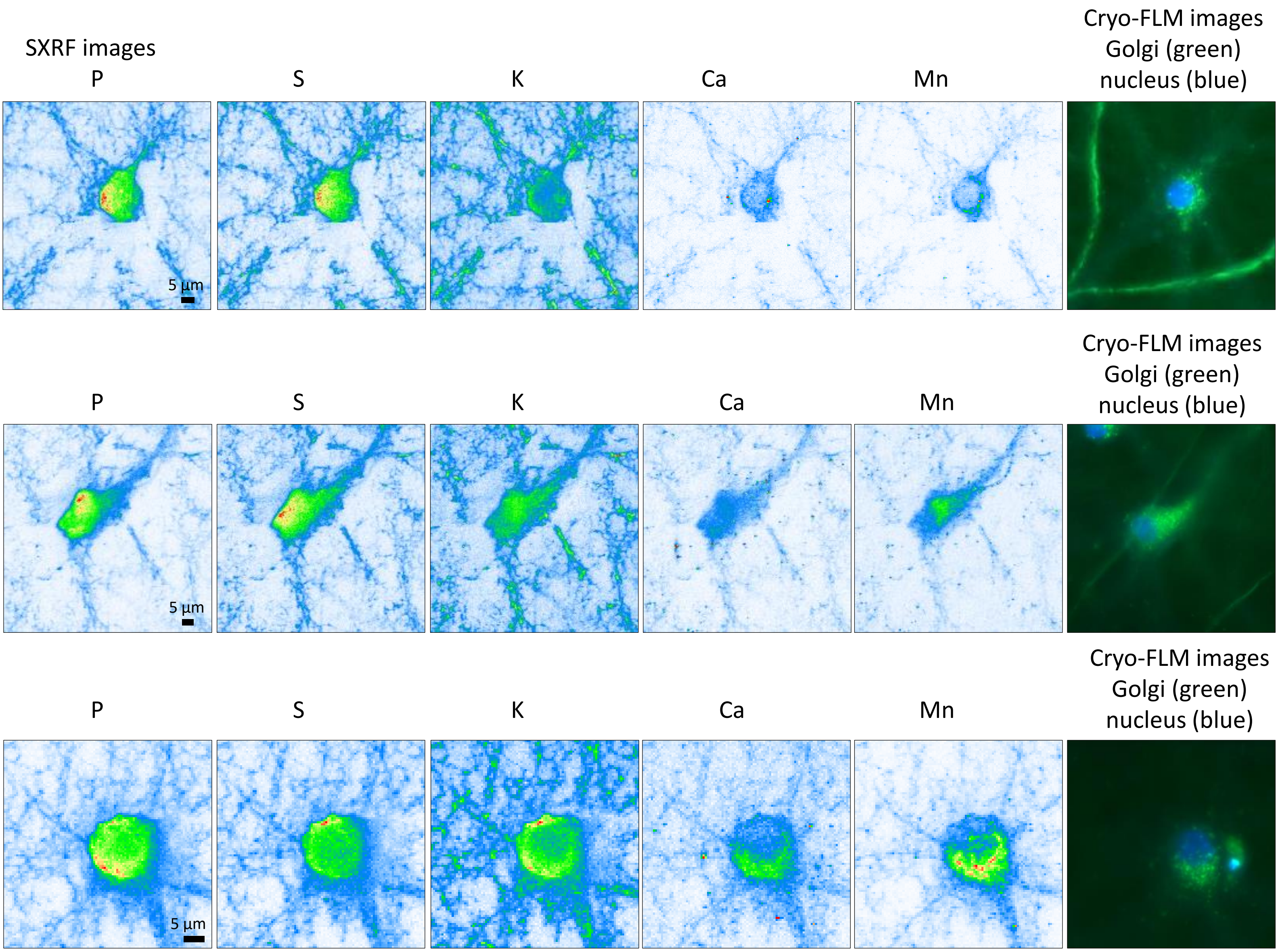

**Supplementary material 11: SXRF imaging in the dendritic network of neurons exposed to manganese.** a) Elemental distribution of P, K, Ca and Mn obtained by SXRF, scan size 45 µm x 60 µm, step 0.15 µm/pixel, scan time 50 ms. b) Merged images of Mn and K evidencing Mn in the neuronal branching, and images of Mn and Ca showing colocalization. c) zoomed region (white square in b) showing Mn along dendrites and Mn and Ca colocalization.

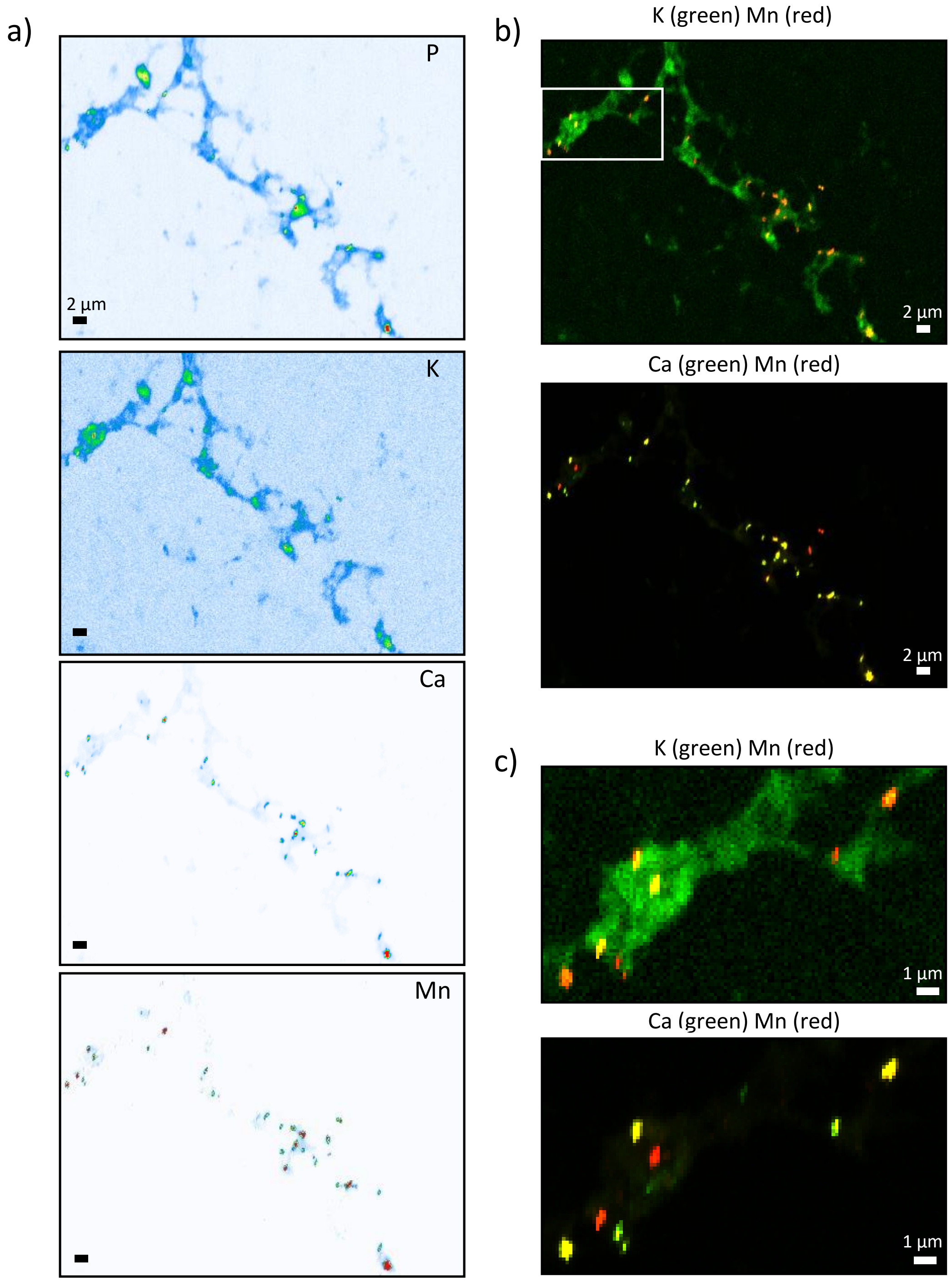

**Supplementary material 12.** Elemental imaging in the dendritic spines of primary neurons exposed to Mn. a) Confocal fluorescence images of tubulin (magenta) and PSD-95 (green) in living primary hippocampal neurons at DIV21. b) Cryo-FLM images of the framed region in a), showing fluorescence of tubulin (magenta) and PSD-95 (green). c) Elemental distribution of Zn and Mn obtained by SXRF in the same region as b), scan size 50 µm x 75 µm, step 0.2 µm/pixel scan time 50 ms/pixel. A 10.2 keV beam was focus to 134 nm x 176 cm, providing a flux of $9.15{\cdot}E^{10}$ photon$\cdot s^{-1}$. d) Superimposed images of Mn (red), PSD-95 (green) and tubulin (magenta) and superimposition of Zn (red), PSD-95 (green) and tubulin (magenta). Zoom in the regions framed in d) to show e) Mn and Zn reaching the post synaptic compartments (blue squares), f) Mn in the close vicinity of dendritic spines (green squares) and g) Mn along the microtubules (red squares). Scale bar 10 µm for panel a; 2 µm for panels b, c and d; 500 nm for panels e, f and g.

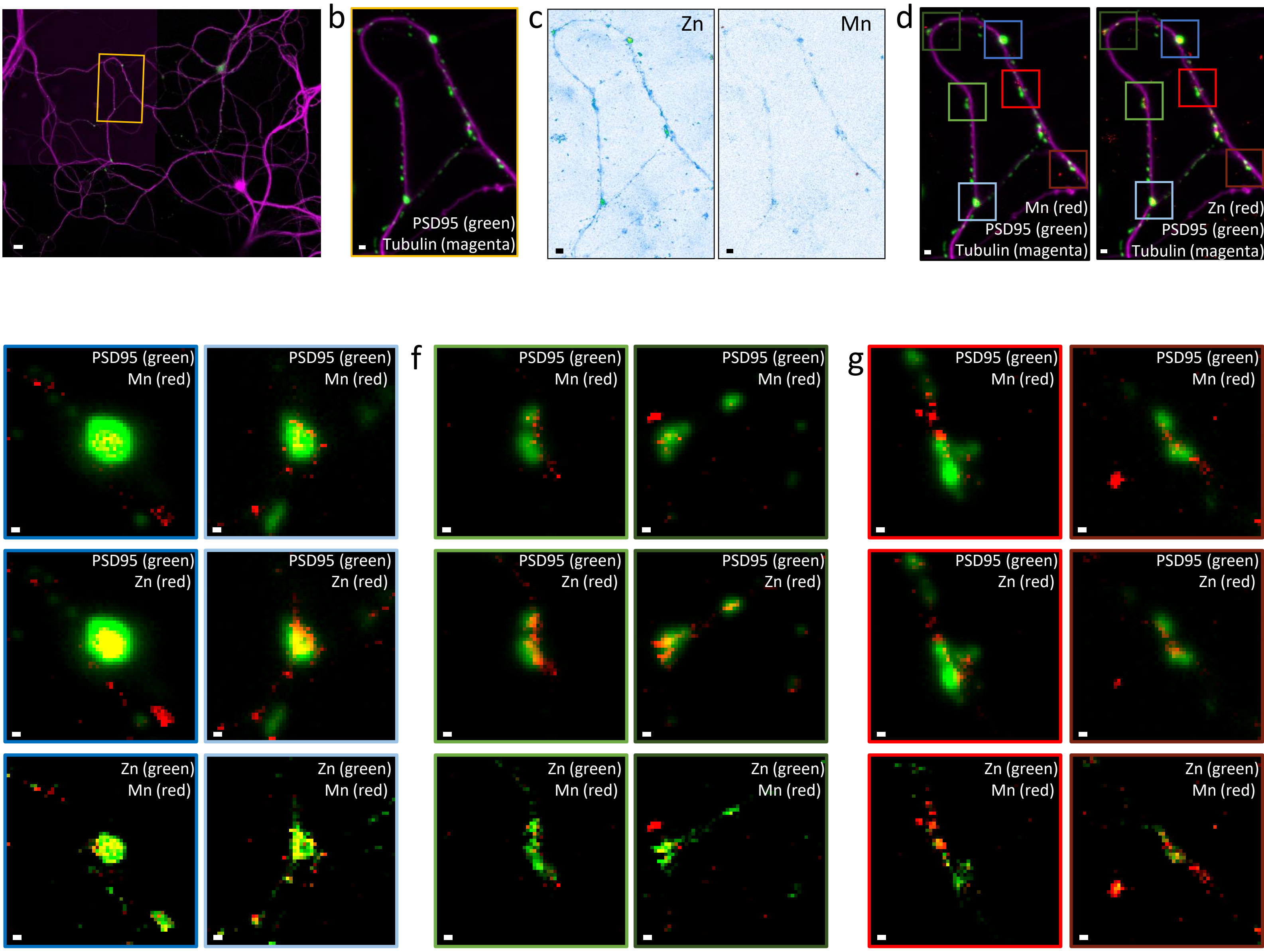
